# High-Dimensional Protein Analysis Uncovers Distinct Immunological and Stromal Signatures Between Primary and Metastatic Pancreatic Ductal Adenocarcinoma

**DOI:** 10.1101/2025.04.18.649381

**Authors:** Emily Greene, Natalie K. Horvat, Deon B. Doxie, Vaunita C. Parihar, Jayden Kim, Cameron J. Herting, Erin E. Grundy, Ayana T. Ruffin, Alyssa M. Krasinskas, Shishir K. Maithel, Juan M. Sarmiento, Mihir M. Shah, Mohammad Y. Zaidi, Maria Diab, Olatunji B. Alese, Kavita Dhodapkar, Haydn T. Kissick, Bassel F. El-Rayes, Chrystal M. Paulos, Gregory B. Lesinski

## Abstract

Our understanding of the pancreatic ductal adenocarcinoma (PDAC) tumor microenvironment (TME) primarily stems from murine models or primary patient tumors. While metastatic tumors have generally less immune infiltration compared to primary tumors, the specific cellular features of metastatic PDAC remain understudied. This knowledge gap is impactful as most patients present with metastatic disease and are most often enrolled in clinical trials. We hypothesized PDAC tumors harbor distinct immunologic and stromal features depending on their anatomical site. Using multiplex immunohistochemistry (mIHC), spatial analysis, and single-cell mass cytometry (CyTOF), we uncover dominant immune and stromal cell populations in tumors derived from 27 primary and 26 liver metastases. Metastatic liver tumors from PDAC patients contained fewer T cells and alpha-smooth muscle actin (α-SMA^+^) activated fibroblasts than primary lesions, while CD68^+^ cells were more abundant. Spatial analyses revealed distinct immune cell communities in primary and metastatic PDAC, whereby CK19^+^ cells clustered differentially with α-SMA^+^, CD3^+^, and CD68^+^ cells, depending on tumor site. When comparing tumor-associated regions, the proportion of peritumoral CK19^-^ cells remained consistent, but their composition varied by disease site. CD8^+^ T cells were significantly less frequent in metastatic tumors, while both CD4^+^ and CD8^+^ T cells present in primary tumors expressed more transcription factors (TFs) associated with suppressive properties, including FoxP3 and ROR*γ*t. CyTOF revealed that T cells co-expressed multiple inhibitory checkpoint receptors, with LAG-3 and PD-1 predominating. This report reveals that primary and metastatic tumors from PDAC patients harbor vastly distinct immunologic and stromal features at the protein level.

**Statement of Significance:** Protein level analysis reveals distinct immunological and stromal features between primary and metastatic PDAC tumors, offering a rationale for immunotherapies that target myeloid cells and increase T cell abundance in metastatic disease.

## INTRODUCTION

Patients with pancreatic ductal adenocarcinoma (PDAC) have a 5-year overall survival of only 13 percent (1). When identified early, resection of primary tumors improves overall survival. However, about 80% of patients recur, usually presenting with metastases (2). The prevailing view is that most patients harbor micro-metastatic disease in the liver or other organs, which is challenging to detect by traditional staging (3). These sobering scenarios provide insight into challenges limiting long-term efficacy of chemotherapy in PDAC patients.

Immunotherapy does not have broad activity in patients with PDAC. This limited efficacy is attributed to multiple factors: 1) poor T cell trafficking, 2) low neoantigen burden, 3) a tumor microenvironment (TME) dominated by suppressive immune populations, and 4) desmoplastic stroma that is a barrier to antitumor immunity. Further complicating matters is that most of the knowledge regarding cellular composition and interactions in human PDAC is derived from primary tumors, rather than from metastases (4-6). We posit this limitation has biased selection of immune targets in clinical trials and may contribute to lack of immunotherapy efficacy in patients with metastatic disease. While clinical efforts have focused on therapeutic combinations with PD-1/PD-L1 blockade, this approach has not yielded advances. Consequently, the role of other inhibitory immune checkpoints or costimulatory molecules remains an active area of interest (7). Emerging data from primary tumors from PDAC patients suggest that alternative ligands such as LAG-3, TIGIT, or TIM-3 may be more abundant targets (8). Despite these efforts, there remains a need to tailor strategies in a data-driven manner, with focus given to immune features in metastatic, rather than primary tumors.

Liver metastases are problematic given their propensity for spread, coupled with the inherent immunosuppressive nature of the liver. Preclinical studies demonstrate liver metastases harbor abundant T regulatory cells (Tregs) (9) and attract activated Fas^+^CD8^+^ T cells from circulation, which undergo apoptosis following interaction with FasL^+^ macrophages (10). Hepatic stellate cells can also facilitate immune suppression, as they promote Tregs and CD11b^+^Gr-1^+^ MDSC differentiation (11). Additionally, studies have documented the propensity of hepatocytes to produce cytokines, such as IL-6, to support a metastatic niche (12). Together these observations, primarily from pre-clinical models, suggest further insight into human PDAC liver metastases is needed to rationally prioritize immunotherapy approaches for future clinical trials.

We hypothesized PDAC tumors harbor distinct immunologic and stromal features depending on where they reside in the body. Using mIHC, we uncovered the dominant immune cell populations within tumors of patients with primary and/or liver metastases. These data indicated a paucity of T cells and alpha-smooth muscle actin^+^ (α-SMA) activated fibroblasts within metastatic, versus primary PDAC. Compared to primary tumors, metastatic PDAC contained a higher density of CD68^+^ macrophages and a modest increase in CD19^+^ B cells. Spatial analyses revealed distinct cellular communities, the composition of which differed between primary and metastatic tumors. More granular analysis of T cells revealed fewer CD8^+^ T cells in metastatic tumors, while CD4^+^ and CD8^+^ T cells in primary tumors expressed more ROR*γ*t and FoxP3 transcription factors as compared to metastatic samples. Single-cell mass cytometry (CyTOF) validated these trends for B and T cells and was used to interrogate more precise phenotypic features of individual cell populations in patient tumors. Finally, both B and T cells co-expressed multiple inhibitory checkpoints, suggesting their functions could be compromised. To our knowledge, these data represent the first robust evaluation of immune and stromal features between primary and metastatic PDAC in patients at the protein level and can be used in prioritizing future immunotherapy strategies in the metastatic setting.

## MATERIALS AND METHODS

Detailed methods are provided in **Supplemental Materials and Methods**.

## RESULTS

### Primary and metastatic PDAC harbor distinct cellular compositions

We posited cellular features of primary and metastatic PDAC tumors in patients differ substantially. To test this concept, we interrogated the composition of primary resected PDAC tumors from consented patients (n=27) and from image-guided, core needle pre-treatment biopsies of patients with liver metastases who were on a clinical trial (n=26) (**Figure 1**). To minimize potential artifacts introduced by the process of surgical resection, we ensured that all samples underwent rigorous quality control measures prior to analysis. Specifically, we evaluated tissue viability and cellular composition to reduce the likelihood that sampling differences significantly impacted our findings. Patient demographics, treatment history, and anatomic site data are in **Supplemental Tables 1 and 2**. All patients received prior therapy, typically a chemotherapy-containing regimen and completed systemic therapy at least two weeks before resection or liver biopsy. Twenty-four primary PDAC tumors and 21 metastases were of sufficient quality for mIHC analysis (**Supplemental Tables 3 and 4**). A portion of fresh tissue was digested into a single-cell suspension and underwent CyTOF analysis, which yielded samples of sufficient numbers of cells for subsequent analysis in 12 primary and 23 metastatic tumors (**Supplemental Table 3 and 4**). Two primary and one metastatic sample were excluded from CyTOF analysis following data capture due to insufficient cell viability.

**Figure 1.**
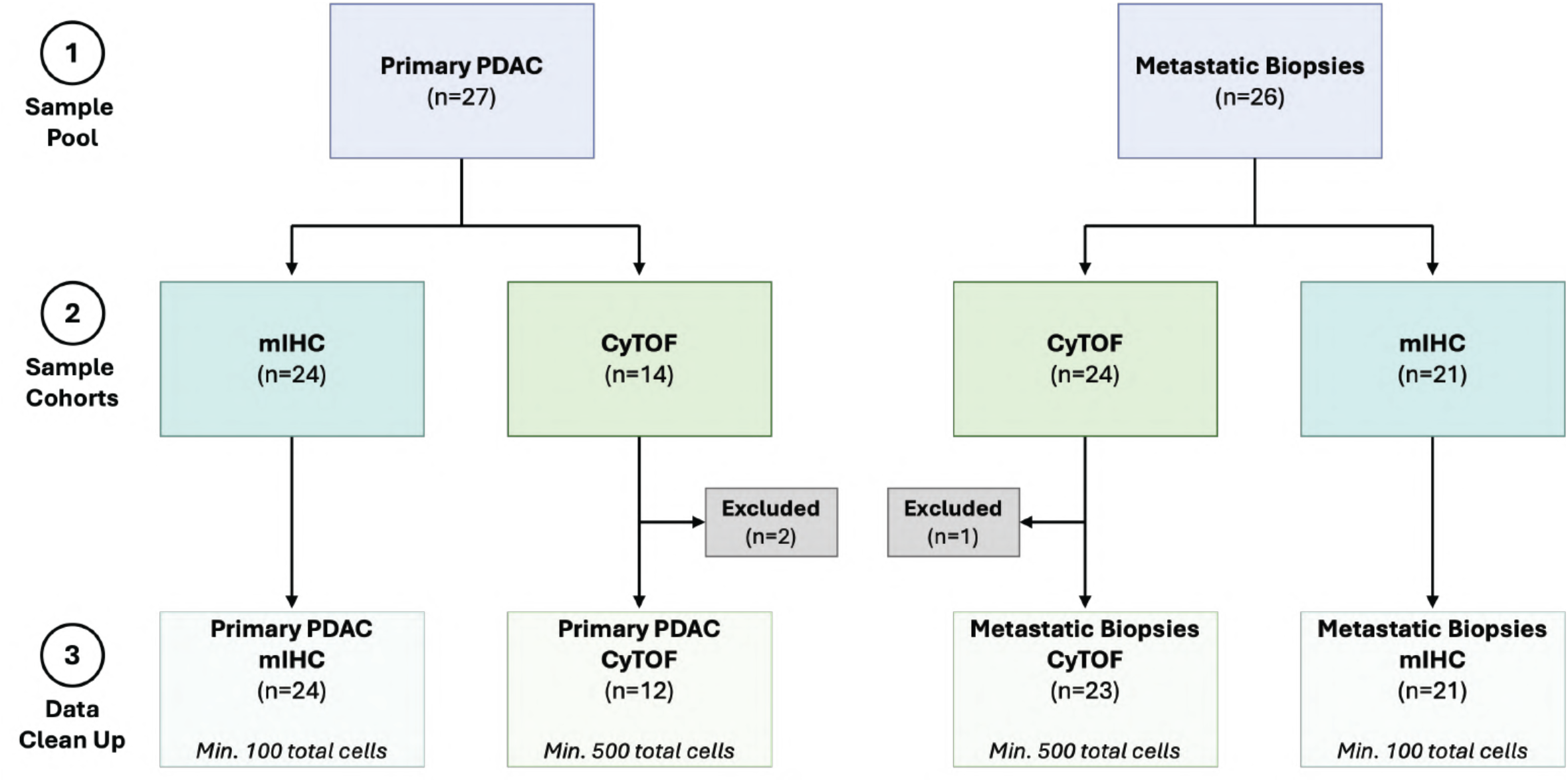
Patient sample workflow: acquisition, processing, and data analysis. Flow chart illustrating (1) total number of samples acquired per tumor type, (2) the number of samples processed via mIHC or CyTOF, and (3) the final number of samples used for data analysis in this study, based on minimum cell number criteria.

Multiplex IHC analysis revealed distinct cell compositions between primary and metastatic tumors (**Figure 2A, Supplemental Figure 1** and **Supplemental Table 5**). Notably, the immune contexture of primary and metastatic tumors was strikingly different (**Figure 2B-D**). For instance, metastatic tumors were nearly devoid of T cells (**Figure 2B and 2D**). Although detectable, these cells comprised only a small fraction of those present within primary tumors. In contrast, CD68^+^ macrophages made up a large proportion of cells in metastatic tumors (primary, 2.44% ± 1.72 vs. 8.29% ± 10.4) (**Figure 2D**). Interestingly, CD19^+^ B cells were present in a subset of metastases but trended toward lower prevalence in primary tumors (**Figure 2D**). Since a salient feature of PDAC is a desmoplastic stroma populated by cancer-associated fibroblasts (CAFs), we quantified α-SMA^+^ cell frequency in the TME, indicative of activated CAFs. These cells were more frequent in primary versus metastatic tumors (25.5% ± 11.3 vs. 11.2% ± 10.6 total nucleated cells, respectively) (**Figure 2B-C**). Subclassification of CAFs using mIHC data, which identified myofibroblast-like CAFs (myCAF; α-SMA^+^IL-6^-^) and inflammatory CAFs (iCAF; α-SMA^+^IL-6^+^) (**Figure 2E**), revealed predominant myCAFs at both sites. However, both myCAF and iCAF were more infrequent in metastatic tumor biopsies (**Figure 2F**). On a per-cell basis, however, the ratio of myCAF to iCAF was more elevated in metastatic compared to primary lesions (**Figure 2G**). These findings were confirmed by evaluating cell density of each population based on tissue area (**Supplemental Figure 2A-H**). Finally, we assessed IL-6 expression across all cells and observed a high propensity of IL-6 expression across tumor sites (**Supplemental Figure 2I-J**), suggesting widespread immune suppressive features within these tumors.

**Figure 2.**
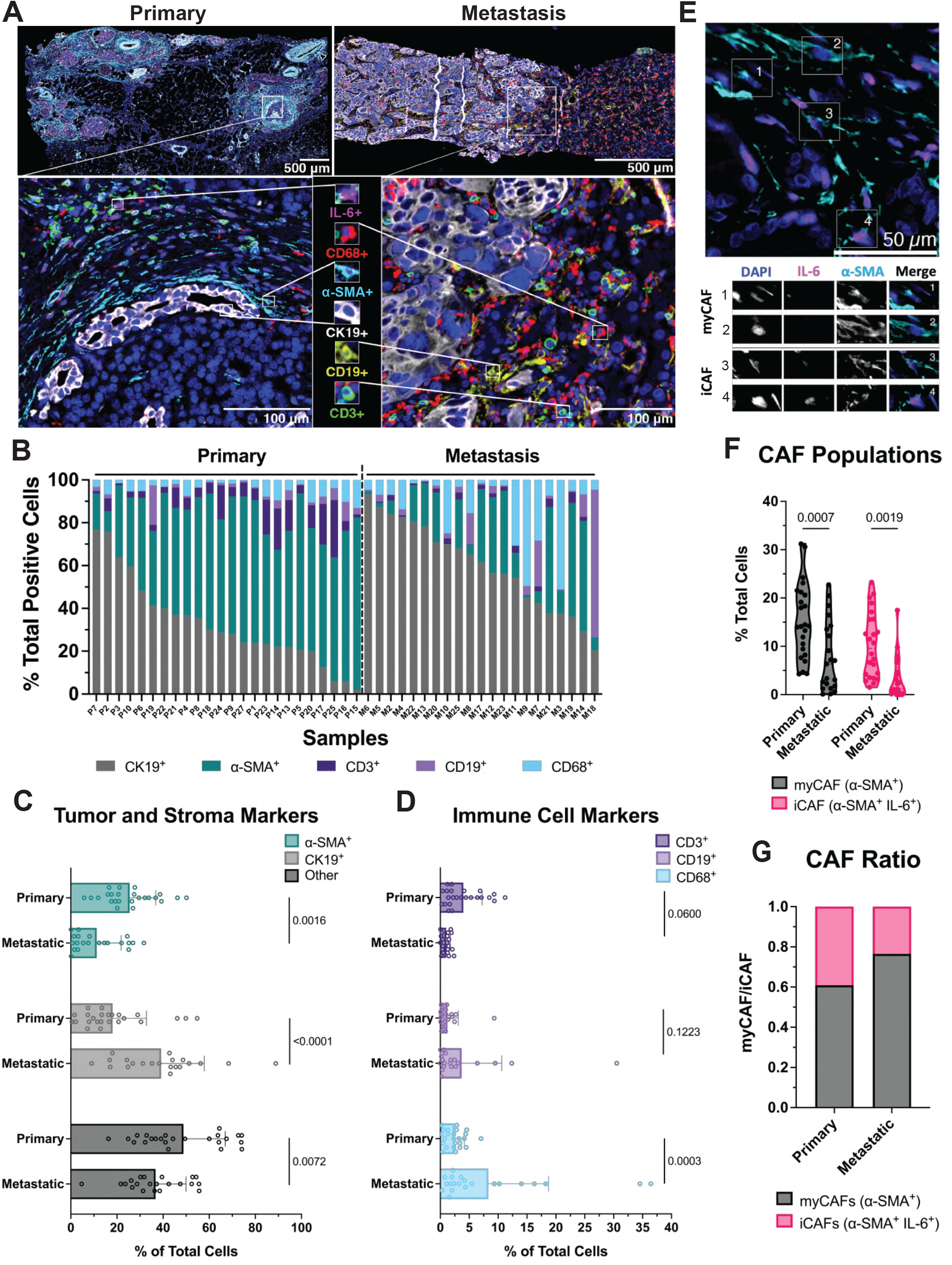
Primary and metastatic PDAC tumors have distinct cellular compositions. (A) Representative fluorescent mIHC images of primary and metastatic tumors. Antibody panel detecting IL-6 (pink), CD68 (red), α-SMA (cyan), CK19 (white), CD19 (yellow) and CD3 (green) indicated by representative cells. (B) Stacked bar graph showing the distribution of CK19^+^, α-SMA^+^, CD3^+^, CD19^+^, and CD68^+^ cells detected by mIHC in each patient sample. Samples arranged from high to low CK19^+^. Bar graphs quantify each detected cell subsets separated into (C) tumor and stroma markers and (D) immune cell markers, shown as a percentage of total cells (Two-Way ANOVA). (E) Representative image of myCAF and iCAF populations in stromal dense tissue (top) with selected CAFs highlighted (myCAF labeled 1 and 2; iCAF labeled 3 and 4) by single channel DAPI, IL-6, and α-SMA images (bottom). (F) Truncated violin plot quantifying myCAF and iCAF populations as a percentage of total cells (Two-Way ANOVA). (G) Stacked bar graph indicating ratio of CAF populations in tumor tissue. Sample size: n=24, primary; n=21, metastatic. PDAC, pancreatic ductal adenocarcinoma; mIHC, multiplex immunohistochemistry; IL-6, interleukin 6; α-SMA, alpha smooth muscle actin; CK19, Cytokeratin 19; CAF, cancer-associated fibroblast; myCAF, myofibroblast; iCAF, inflammatory fibroblast.

### Differences in immune cell distribution and proximity to CK19^+^ cells between primary and metastatic PDAC

To investigate distribution and organization of cells in primary and metastatic PDAC, *x* and *y* coordinates of phenotypically defined cell populations extracted from whole tissue images (**Figure 2**) were mapped post-mIHC for all samples that passed quality control metrics. Median and minimum distances between all cell types were measured in both primary (n=24) and metastatic (n=20) tissues, showing little difference in cell dispersal and closeness between the two tissues (**Supplemental Figure 3A-B**). To focus on cellular interactions, minimum distance (μm) measurements between phenotyped cells were obtained (**Figure 3A** and **Supplemental Table 6**), and results were summarized in a heatmap with hierarchical clustering of target cells. We found that αSMA^+^, CD19^+^IL-6^+^, and CD68^+^ cells were located closer to CK19^+^ cells in metastatic PDAC than primary tumors (**Supplemental Figure 3C**). Consistent with other studies and phenotypic properties of iCAFs (13), we also found that αSMA^+^IL-6^+^ cells were distant from primary tumors, but in closer proximity to CK19^+^ cells in metastases. In primary PDAC, IL-6^+^ cells were positioned farther from CK19^+^ cells than IL-6^-^ cells. In metastatic tissue, only CD3^+^IL-6^+^ cells exhibited greater distance from CK19^+^ cells (**Figure 3A** and **Supplemental Table 6**). Given our data suggesting more abundant CD68^+^ and CD19^+^ cells in metastatic lesions, we focused on spatial relationships around each as the reference cell types. These data indicated that CD19^+^ cells reside closer to CK19^+^ cells in metastatic tissue (**Supplemental Figure 3D**). Across both sites, CD19^+^ cells were most closely associated with CD68^+^ and CD3^+^ cells but remained distant from α-SMA^+^ and IL-6 co-expressing cells. In metastatic PDAC, CD68^+^ cells were positioned closer to CK19^+^ and α-SMA^+^ cells. At both sites, CD68^+^ cells closely localized with CK19^+^ and CD3^+^ cells (**Supplemental Figure 3E**).

**Figure 3.**
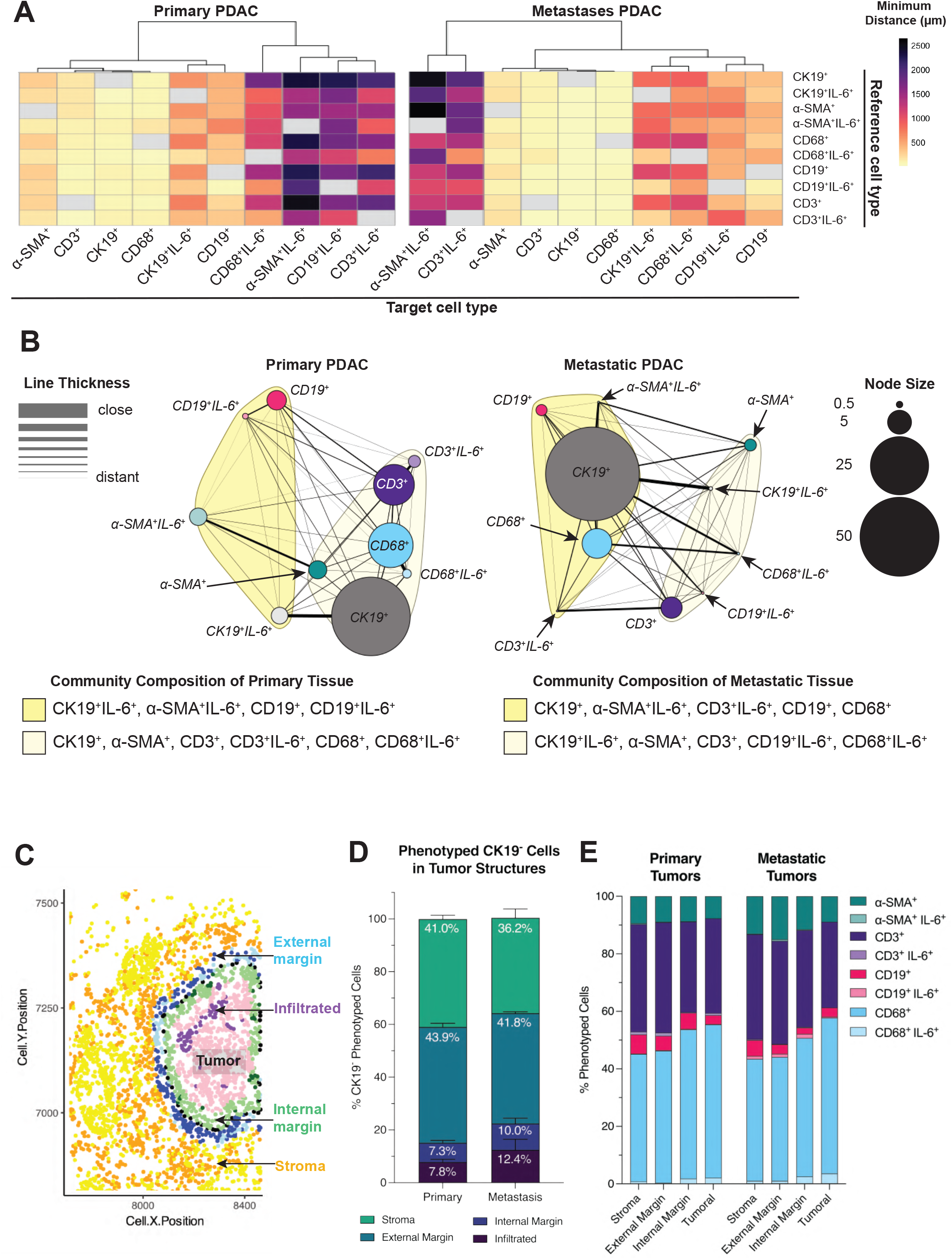
Cellular organization around CK19^+^ cells exhibit marked differences between primary and metastatic PDAC. (A) Minimum distances measured (μm) between reference cell (y axis) and target cell (x axis) in primary and metastatic PDAC tissue represented as heatmaps with hierarchical clustering of target cell types. (B) Network diagram of primary and metastatic PDAC tissue with percentage of cell type represented as nodes size and distances between cell types represented as line thickness. Cloud coloring around node groups represents communities. (C) Representative dot plot depicts four levels of tumor structures: stroma, external margin, internal margin, infiltrated into the tumoral region. Stacked bar graphs indicating the percent of (D) total CK19^-^ phenotyped cells and (E) identified cell subsets in each tumor structure of primary and metastatic PDAC tissue (Two-Way ANOVA). Sample sizes: tissue spatial analysis: n=24, primary; n=20, metastatic; tumor structure analysis: n=20, primary; n=10, metastatic. CK19, Cytokeratin 19; PDAC, pancreatic ductal adenocarcinoma.

To summarize cell dispersion patterns in primary and metastatic tumors, directional cell-to-cell distances and percentages were clustered and visualized as a network diagram, with close cells identified as communities (**Figure 3B**). Communities containing CK19^+^ cells varied between tumor types. In primary PDAC, CK19^+^ epithelial cells clustered with α-SMA^+^ fibroblasts, CD3^+^ T cells, CD3^+^IL-6^+^, CD68^+^ macrophages, and CD68^+^IL-6^+^ cells. In contrast, metastatic PDAC showed CK19^+^ cells grouping with α-SMA^+^IL-6^+^, CD3^+^IL-6^+^, CD19^+^ B cells, and CD68^+^ cells— suggesting a shift in cancer–immune interactions during metastasis. Notably, in primary tumors, CD19^+^ B cells were linked to CK19^+^IL-6^+^, α-SMA^+^IL-6^+^, and CD19^+^IL-6^+^ cells, whereas in metastases, a distinct cellular community emerged, composed of CK19^+^IL-6^+^, α-SMA^+^, CD3^+^, CD19^+^IL-6^+^, and CD68^+^IL-6^+^ cells.

Tissue architecture was assessed by the ratio of peritumoral cells (CK19^-^) to intratumoral cells (CK19^+^), excluding samples with ambiguous margins from further analysis. This method mapped distinct regions, such as intratumoral, internal, external margins, and stroma, across twenty primary and ten metastatic tissues (**Figure 3C**). Quantification of CK19^-^ phenotype cells revealed similar fractions within each structure (**Figure 3D, Supplemental Figure 3F** and **Supplemental Table 7**). In-depth analysis of cell subsets relative to mapped tumor regions (**Figure 3E** and **Supplemental Table 8**) identified a predominance of CD3^+^ T cells in stromal and external margins in both primary and metastatic tumor areas (**Supplemental Figure 3G-J**), suggesting that T cells are less likely to penetrate the tumor. Notably, CD68^+^ cells were more abundant in internal margins and tumoral regions across sites (**Supplemental Figure 3G-J**). Moreover, metastases contained more α-SMA^+^ cells around the external margin (**Supplemental Figure 3G-J**). The overall presence of CD19^+^ B cells across tumor regions did not differ between disease site (**Supplemental Figure 3G-J**) suggesting that CD19^+^ B cells are prevalent throughout cancerous tissue.

### T cells from primary PDAC tumors display prominent expression of FoxP3 and ROR*γ*t

T cell subsets were more rigorously evaluated using mIHC (**Figure 4A-B, Supplemental Figure 1** and **Supplemental Table 5**). Consistent with data using the pan-T cell marker, CD3, very few CD8^+^ T cells were detected, with significantly greater frequency in primary as compared to metastatic tumors (7.99% ± 8.95 or 0.90% ± 0.91 total nucleated cells, respectively, **Figure 4C-D**). The limited CD8^+^ T cells in metastases precluded more detailed analysis of transcription factors, or the small populations of T cells co-expressing CD4 and CD8 (**Figure 4C-D**). CD4^+^ T cells were present at greater frequency in primary (3.19% ± 4.16 total nucleated cells, **Figure 4D**) and metastatic (5.04% ± 5.79 total nucleated cells, **Figure 4D**) PDAC tumors, enabling characterization of transcription factors (TF) with greater confidence. We noted that FoxP3 and/or ROR*γ*t TFs were predominant in CD4^+^ T cells from primary tumors compared to metastatic tumors (**Figure 4E-F**). A trend towards increased Th1-promoting T-bet expression in metastatic samples may suggest a more inflammatory phenotype in these cells. While CD8^+^ T cells were fewer overall in tumors, T-bet expression was also predominant, regardless of tumor sites. Although the overall frequency was quite limited, a fraction of these T cell populations did also express FoxP3 and ROR*γ*t (**Supplemental Figure 4**),

**Figure 4.**
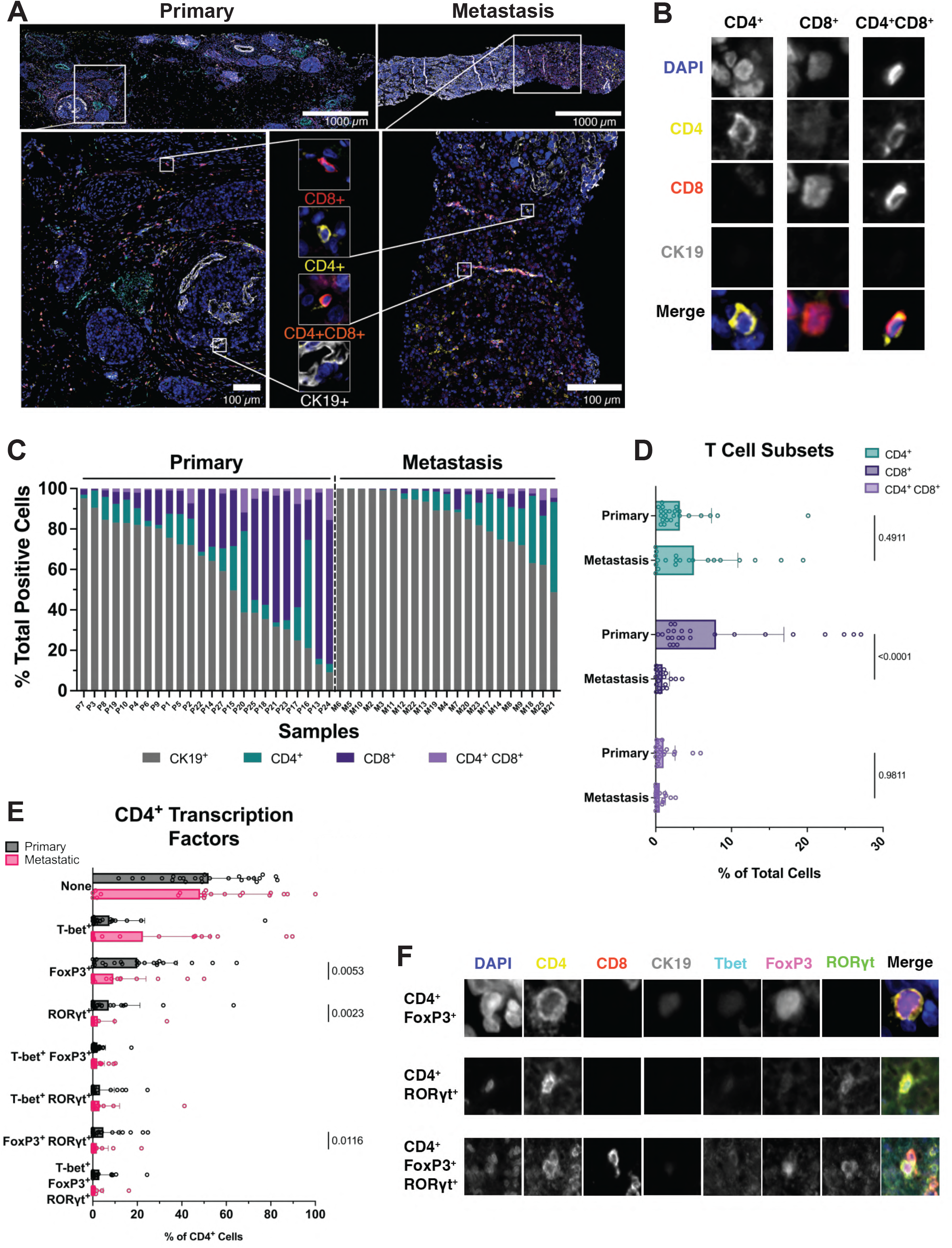
T cells in PDAC tumors are limited and express transcription factors with suppressive phenotypic properties. (A) Representative images of mIHC staining on primary and metastatic tumor samples. Antibody panel detecting CD8 (red), CD4 (yellow), CK19 (white), and dual stained CD4^+^CD8^+^ cells (orange), indicated by representative cells. (B) Representative images of CD4^+^, CD8^+^ and CD4^+^CD8^+^ cell subsets by single channel DAPI, CD4, CD8 and CK19 images. (C) Stacked bar graph showing distribution of CK19^+^, CD4^+^, CD8^+^ and CD4^+^CD8^+^ cell detected by mIHC in each patient sample. Samples arranged from high to low CK19^+^. (D) Bar graphs showing the mean and SD of the percentage of each measured T cell subset (Two-Way ANOVA). (E) Bar graph showing the mean and SD of the percentage of transcription factor (T-bet, FoxP3 and ROR*γ*t) positive CD4^+^ T cells populations represented as a total of CD4^+^ T cells (Mann-Whitney tests). (F) Representative images of CD4^+^ T cell subsets by single channel DAPI, CD4, CD8, CK19, T-bet, FoxP3 and ROR*γ*t images. Sample size: n=24, primary; n=21, metastatic. PDAC, pancreatic ductal adenocarcinoma; mIHC, multiplex immunohistochemistry; CK19, Cytokeratin 19; SD, standard deviation.

### CyTOF analysis reveals immunoregulatory features of lymphocyte populations residing in primary and liver metastatic PDAC

Mass cytometry is advantageous for providing single-cell resolution from small tissues. Given the overall paucity of T and B cells within PDAC tumors, we used this technology and employed a tissue digestion protocol optimized for lymphocyte preservation. A CyTOF panel enriched with antibodies against lymphocyte markers was used to interrogate cellular features with greater scrutiny at the single cell level (**Supplemental Table 9)**. For these assays, freshly obtained patient tissues were processed to obtain single-cell suspensions. Following quality control, our cohort for analysis consisted of n=12 primary and n=23 metastatic tumors. Clustering data for immune lineages from concatenated data are in **Figure 5A**. There was noticeable heterogeneity, even amongst samples derived from the same tumor site. For example, primary samples P25 and P9 contained almost no immune cells, while CD45^+^ infiltration was high in samples P7 and P27 (**Figure 5B**). Similarly, while most cells in metastases were CD45^-^, samples such as M8 and M9 harbored detectable CD45^+^ cells expressing CD4, CD8 and/or CD19 indicative of lymphocytes (**Figure 5C**). Consistent with mIHC, primary tumors, on average, had more immune cells than metastases (36.23 ± 35.25% and 33.25 ± 36.79% CD45^+^ cells, respectively; **Supplemental Figure 5A** and **Supplemental Table 10**). Evaluation of lymphocyte lineage showed CD8^+^ T Cells and CD19^+^ B cells trended higher in metastatic versus primary tumors. Consistent with these data, CD8^+^ T cells and CD19^+^ B cells within metastases had more frequent positivity for the Ki-67 proliferation marker (**Supplemental Figure 5B** and **Supplemental Table 10**). Enzymatically processing tissue reduced cell frequency across samples, limiting in-depth analysis of additional lineage markers and transcription factors; however, exploratory analyses were performed where possible (**Supplemental Figure 6-10** and **Supplemental Tables 10-12**).

**Figure 5.**
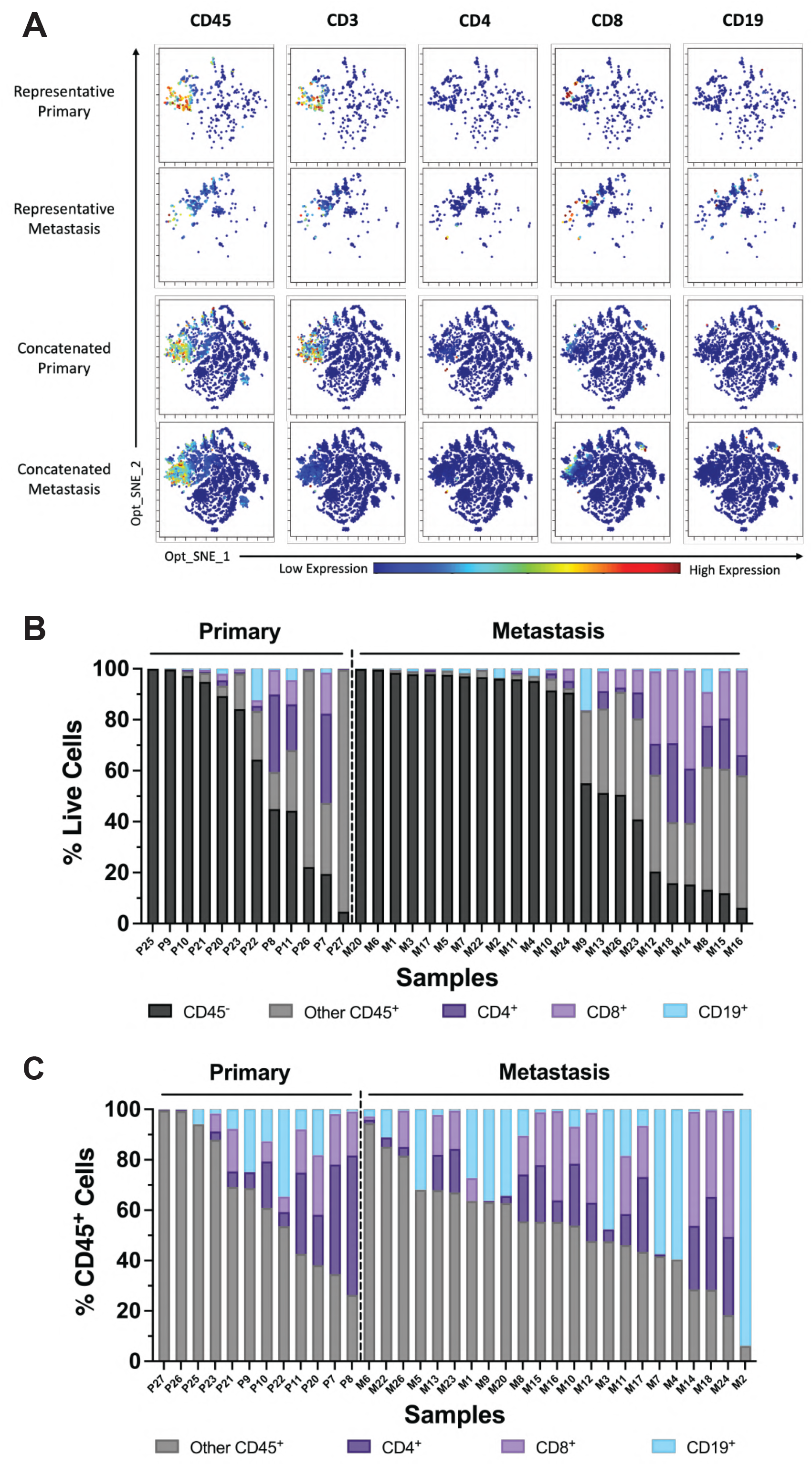
Single-cell analysis of lymphocyte populations by mass cytometry validates reduced T cells and increased B cells in liver metastases. (A) Levels of CD45, CD3, CD4, CD8 and CD19 are displayed on individual, unsupervised Opt_SNE plots of live single cells in representative (top) and concatenated (bottom) primary and metastatic samples. (B) Stacked bar graph of the distribution of CD45^-^, CD45^+^, CD4^+^, CD8^+^ and CD19^+^ cells shown as a percentage of total live cells detected in each patient by mass cytometry. Samples arranged from high to low CD45^-^. (C) Stacked bar graph of the distribution of CD45^+^, CD4^+^, CD8^+^ and CD19^+^ shown as a percentage of total CD45^+^ cells detected in each patient by mass cytometry. Samples arranged from high to low other CD45^+^. Sample size: n=12, primary; n=23, metastatic. Opt_SNE, optimizes t-distributed stochastic neighbor embedding.

CD19^+^ B cells were evident in PDAC tumors, particularly the metastases. To characterize their phenotype, we analyzed tumor infiltrating lymphocytes (TIL) from patients using CyTOF. For robust analysis, a cutoff of 25 CD19^+^ events was applied. Six different phenotypically defined B cell subsets were evaluated (**Supplemental Figure 11A, Supplemental Figure 12** and **Supplemental Tables 13**), revealing several key findings: First, there was a trend toward a higher frequency of naïve B cells in primary tumors versus metastases. Although naïve B cells were present in primary tumors, metastatic tumors showed a trend toward a greater proportion of B cells with regulatory potential, as indicated by PD-1 and/or PD-L1 expression. Also, a small population of CD27^+^ B cells, a marker of memory phenotype, was detectable in a subset of tumors, though their frequency did not differ between primary and metastatic sites. B cells with extrafollicular or plasmablast phenotypes were rare across all samples. Lastly, we accessed activation markers on B cell subpopulations with regulatory potential, which showed high expression levels across both primary and metastatic tumor sites (**Supplemental Figure 11B-C** and **Supplemental Figure 12**).

### T cells express inhibitory immune checkpoint receptors across disease sites

We next evaluated expression of targetable inhibitory checkpoint receptors on CD4^+^ and CD8^+^ T cells in primary and metastatic PDAC. These checkpoints included CTLA-4, LAG-3, PD-1, TIGIT and TIM-3 (**Supplemental Figure 13-14**). We applied a 25-cell cutoff to CD3^+^ T cells for robust data (**Supplemental Table 11**). This resulted in 8 primary and 13 metastatic samples with sufficient events for comparison (**Figure 6A**). Both CD4^+^ and CD8^+^ T cells from patients co-express multiple inhibitory checkpoints (**Figure 6B-C**). Further analysis indicated CTLA-4, LAG-3 and PD-1 were expressed most abundantly on CD4^+^ T cells both in primary and metastatic (**Figure 6D**). A trend toward higher TIGIT and TIM-3 on CD4^+^ T cells was evident in primary versus metastatic tumors (41.26% vs. 38.38% of TIGIT, respectively and 31.98% vs. 28.54%, respectively). Notably, checkpoints on CD8^+^ T cells were more variable (**Figure 6F**). For example, CTLA-4 and TIGIT were more highly expressed in metastasis than primary samples (60.07% vs. 42.06% of CTLA-4 and 62.84% vs. 50.61% of TIGIT, respectively). Boolean gating of these five markers was performed to assess frequency of concurrent checkpoint expression combinations (**Figure 6E and 6G**). In primary tumors, CTLA-4, LAG-3 and PD-1 co-expression, followed by CTLA-4, LAG-3, PD-1, and TIGIT co-expression were dominant, encompassing >30% of CD4^+^ T cells. In metastases, these same checkpoint combinations dominated on >35% of CD4^+^ T cells (**Figure 6E** and **Supplemental Tables 14**). On CD8^+^ T cells, co-expression of LAG-3 and PD-1, followed by CTLA-4, PD-1 and TIGIT were most prevalent in primary (encompassing >20%), while co-expression of CTLA-4, PD-1 and TIGIT, followed by PD-1 and TIGIT were prevalent in metastatic samples (encompassing >20%) (**Figure 6G** and **Supplemental Tables 15**).

**Figure 6.**
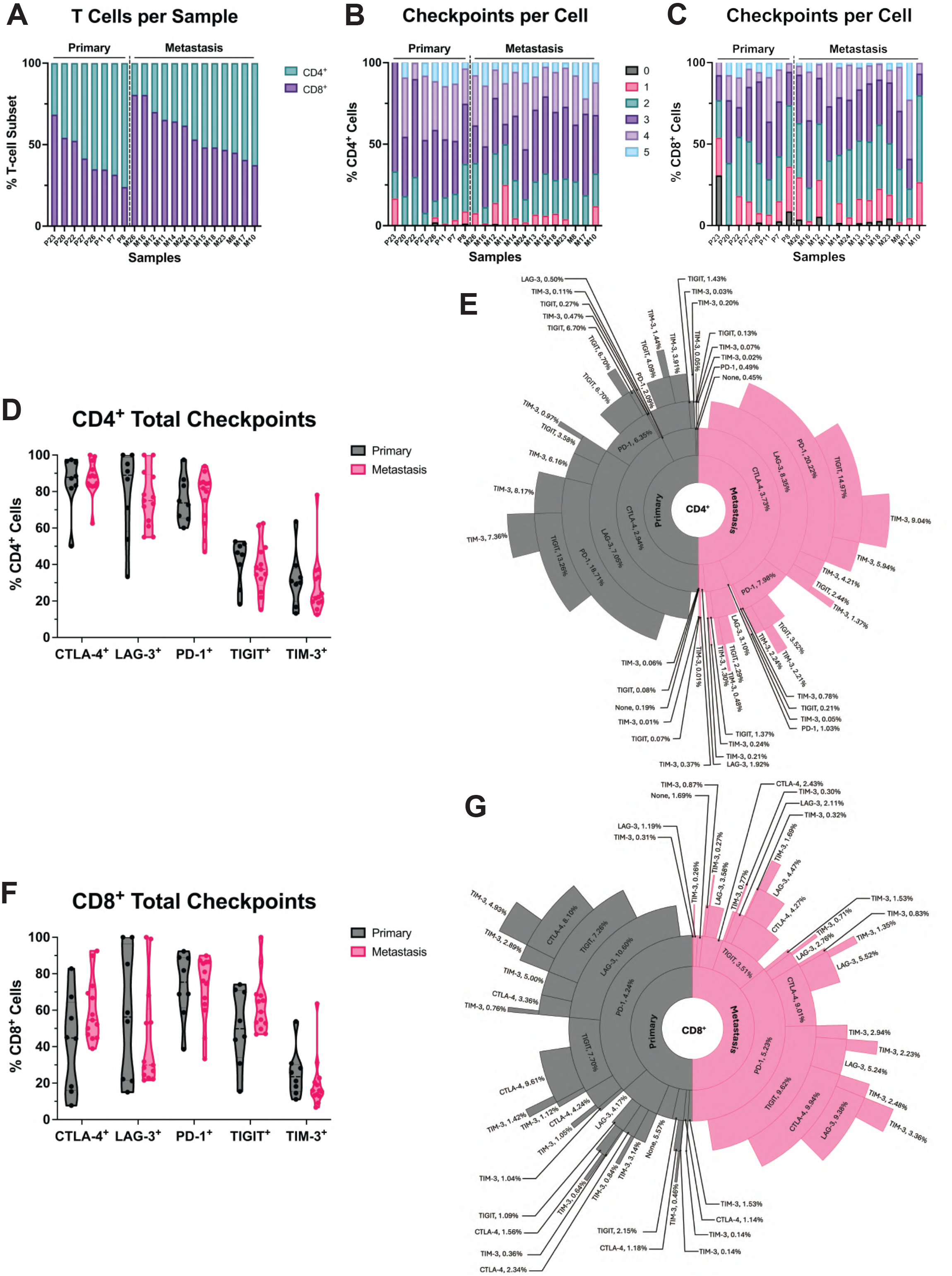
PDAC-associated T cells express prominent inhibitory immune checkpoint receptors regardless of disease site. (A) Stacked bar graph of CD4^+^ and CD8^+^ T cells shown as percentage of total T cells per sample. Samples arranged from high to low CD8^+^ T cells. Stacked bar graphs showing the distribution of checkpoint proteins detected on (B) CD4^+^ T cells and (C) CD8^+^ T cells. Samples arranged in the same order as figure A. Truncated violin plots quantifying CTLA-4, LAG-3, PD-1, TIGIT and TIM-3 checkpoint marker expression on (D) CD4^+^ T cells and (F) CD8^+^ T cells. Sunburst charts of checkpoint marker combinations of CTLA-4, LAG-3, PD-1, TIGIT and TIM-3 found for (E) CD4^+^ and (G) CD8^+^ T cells with the percentage represented for the terminal checkpoint in each combination. Sample size: n=8 primary; n=13 metastatic.

### Multiplex IHC analysis of matched primary and metastases validates trends of larger cohort analysis

Although remarkably rare and challenging to obtain in practice, we acquired matched primary and metastatic PDAC tissues from four patients with available archival specimens (**Supplemental Figure 15A**). Although these tissues were included as part of the larger cohort, this separate analysis of matched patient tissues enabled a direct comparison of trends in TME changes by PDAC location. Consistent with findings from our unmatched patient cohort (**Figure 2**), patient-matched metastases exhibited trends towards higher proportions of CK19^+^ and CD19^+^, with CD68^+^ cells consistently elevated in metastasis of all four patients analyzed (**Supplemental Figure 15B-C**). In contrast, α-SMA^+^ and CD3^+^ cells were more abundant in primary tumors (**Supplemental Figure 15B-C**). Additionally, metastases had more myCAFs and fewer iCAFs, when compared to matched primary tumors in three of four patients (**Supplemental Figure 15D**). In these samples, CD8^+^ T cells trended to decrease between matched primary and metastatic PDAC while there was no trend for CD4^+^ T cells (**Supplemental Figure 15E-F**).

## DISCUSSION

This study uncovers crucial differences in cellular composition of the TME between primary and metastatic PDAC, as visually summarized in **Figure 7**. While a handful of elegant studies have compared either normal and cancerous tissue (8) or profiled primary and metastatic PDAC using single-cell transcriptomics (14-18), our study is the first to report high dimensional protein-level data across disease sites. Given how rare and difficult it is to obtain metastatic PDAC biopsies from PDAC patients for research, the observations from these samples are particularly valuable and informative.

**Figure 7.**
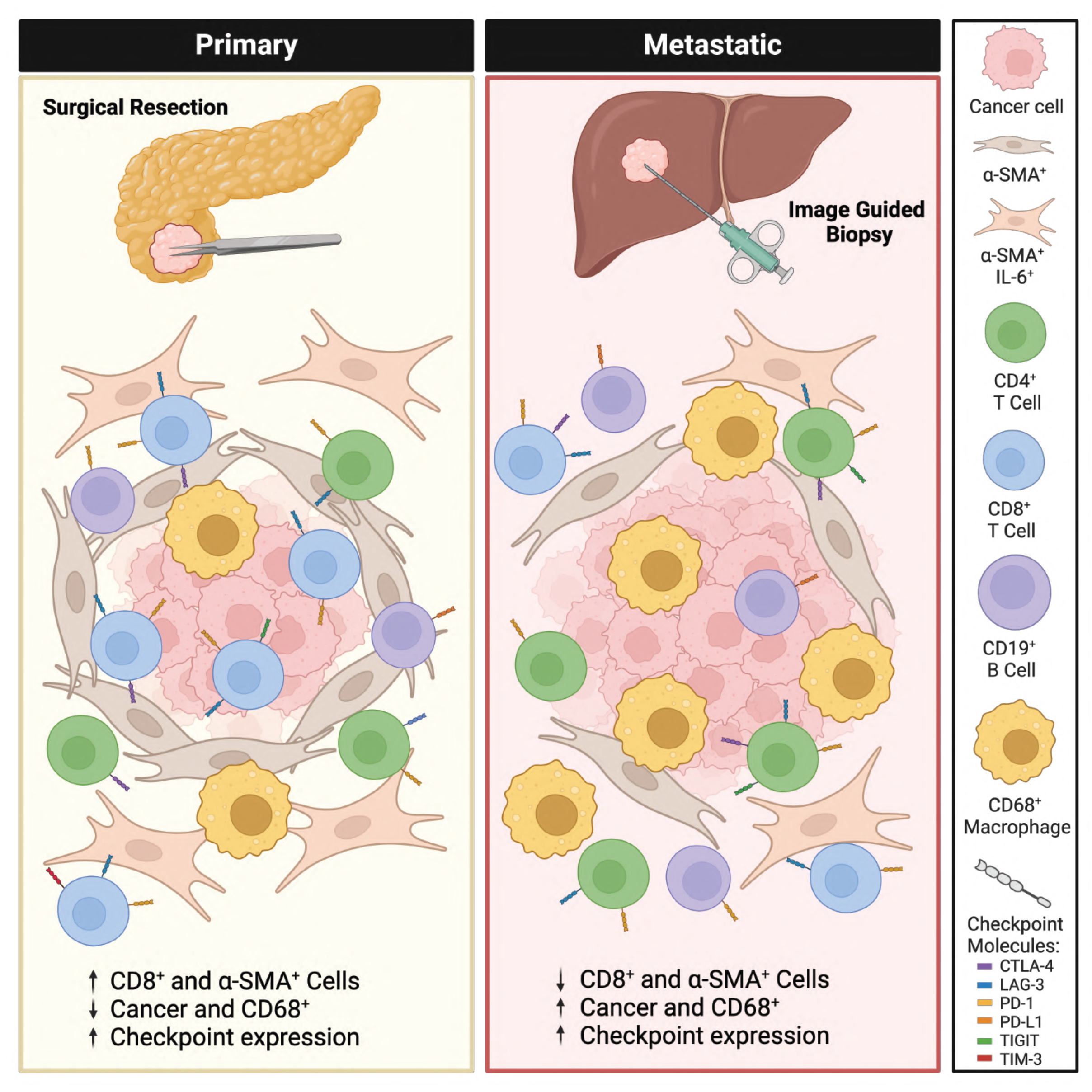
Summary figure depicting differences in stromal and immune cell composition between primary and metastatic PDAC. α-SMA^+^ CAFs and T cells are less prevalent in metastatic tumors, while CK19^+^, CD19^+^ and CD68^+^ cells are more frequent. Spatial analysis reveals cellular neighborhoods are different based on tumor site. Checkpoint marker levels remain high, regardless of tumor location, but the prevalence of each checkpoint and the dominant combinations differ between primary and metastatic tissue. Created in BioRender. Lesinski, G. (2025) https://BioRender.com/w97c106

It is important to note that most prior transcriptomic studies analyzed smaller patient cohorts and lacked protein-level validation. By contrast, our study leverages multiplexed imaging and flow cytometry to characterize lineage-specific markers across a broader set of patient tissues, offering a critical and complementary perspective to the existing literature. While our protein-based panels include fewer phenotypic markers than transcript-based methodology, they were purposefully designed to capture key immune and stromal populations in the PDAC TME and allowed rigorous confirmation of lineage populations in samples across multiple platforms. Notably, the absence of certain immune types in our dataset, such as low T cell frequencies, is consistent with prior reports focused on primary PDAC (8).

Among our findings, we observed a scarcity of T cells, especially in metastatic tumors. In contrast, CD68^+^ cells were more abundant in metastases, accompanied by a trend towards increased CD19^+^ B cells, indicating these populations may play a unique role in this setting. Conversely, CD8^+^ T cells were less frequent in metastasis sites. Spatial analysis indicated CD19^+^, CD68^+^, and CD3^+^ cells are present as ‘communities’ in both primary and metastatic tumors, and that they localize to margins. In metastatic tumors, CK19^+^ cells associated closely with α-SMA^+^, CD19^+^ IL-6^+^ and CD68^+^ cells. Further analysis revealed CD4^+^ T cells expressed ROR*γ*t and FoxP3 transcription factors, along with inhibitory checkpoint receptors, indicative of a suppressive phenotype. These data provide novel insight into unique cellular features in PDAC by disease site and provide rationale for prioritizing targeted and immunotherapy approaches that restore T cells into the metastatic PDAC TME, such as inhibitors of oncogenic Ras signaling, or even adoptive cell transfer (ACT) therapy.

The concept of whether CAFs promote or restrain PDAC progression has been a topic of controversy and interest. Many assumptions regarding interactions between CAFs and PDAC are from murine models or primary human tumors, with more limited data available on their frequency or role in metastatic disease. Not surprisingly, primary PDAC harbored abundant α-SMA^+^ cells, and consistent with other studies (19), these cells were also present in metastases, albeit less frequently in our cohort of heavily pre-treated patients. We also recognize that the method of acquiring these metastatic lesions via tumor targeted, image guided biopsy may have intentionally yielded regions with less stroma. There was also a trend toward these cells adopting a myCAF (IL-6^-^) phenotype (13). Together, these data suggest strategies to deplete or modify stroma may impact patients less with refractory metastatic disease and may provide insight as to why clinical trials targeting this feature of PDAC tumors have, to date, been underwhelming in patients. While these data are relevant, we acknowledge the inherent heterogeneity and plasticity of CAFs which we were unable to comprehensively assess with a limited panel of mIHC markers, or discern from the contribution of hepatic stellate cells or others of relevance (16). Nonetheless, α-SMA remains a well-characterized marker of activated CAFs on most subtypes and speaks to the utility of this data, even with a somewhat limited phenotypic panel.

Our observation that PDAC metastases harbored a greater proportion of CD68^+^ cells was interesting. Data from primary tumors and murine models suggest that myeloid cells in PDAC represent sought-after targets to reverse suppressive immune features (3,6). While this multiplex IHC panel was focused on CD68 and did not have capacity to accommodate other macrophage markers, our data remains consistent with several other studies reporting the role of macrophages in PDAC (20-22). These data also suggest the reliance of PDAC on myeloid cells is further amplified as tumors progress. We speculate this “myeloid amplification” may be fueled by inflammatory cytokines in liver or skew chemokine gradients to attract them efficiently. Indeed, our prior data with patient-derived CAFs revealed a prominent role for IL-6, SDF-1, GM-CSF, and others that influence myeloid biology (23). These results aligned with other work showing SDF-1 from CAFs could be co-targeted to enhance anti-PD-L1 immunotherapy (24). When we analyzed blood of untreated patients with metastatic PDAC, circulating plasma IL-6 was associated with worse overall survival (25). These data suggest cytokine and chemokine circuits that foster myeloid cells, at the expense of dendritic cells, may further amplify disease progression and represent viable therapeutic targets (26-30). Further, a study by Lee *et. al*. demonstrated IL-6/STAT3 signaling leads to production of serum amyloid A1 and A2 (SAA). This increase in SAA from hepatocytes favored a pro-metastatic niche in the liver (12). The fact that CD68^+^ cells predominated metastatic lesions, and T cells were few, supports other conclusions from work by Yu *et al*. (10). Using pre-clinical models, these studies showed macrophages in liver metastases elicit Fas-FasL mediated apoptosis of CD8^+^ T cells, eliminating them from the TME. Data herein are significant as, to date, macrophage or myeloid-targeted therapeutic approaches, including CSF-1R and CCR2/CCR4-directed therapies, have not made a dramatic impact in metastatic PDAC when combined with other approaches in clinical trials (31-33). Likewise, initial promising clinical data combining anti-PD-1 and chemotherapy with CD40 agonists unfortunately did not show superiority to anti-PD-1 and chemotherapy in a randomized phase 2 trial (34,35). Given these disappointing outcomes with myeloid-targeted agents, further scrutiny of myeloid and macrophage phenotypes in metastatic disease could be informative. For instance, future studies could more precisely define dominant chemokine or cytokine receptors on these cells for optimal targeting.

The trend toward higher frequencies of CD19^+^ B cells in metastatic PDAC tumors was unexpected. While other studies indicate B cells are detectable, limited data on their nature in metastases were available (14,16,17,36). Although our mass cytometry panel was intentionally focused on T cell markers, it did allow for interrogation of some key proteins on B cells in select patients at single-cell resolution. Regardless of site, B cells displayed features associated with immune suppression, rather than a stimulatory phenotype. These data argue B cells in tumors are unlikely to facilitate antigen presentation or productive antitumor responses. The idea that B cells support PDAC has inspired studies to targeting Bruton tyrosine kinase (BTK) to interfere with suppressive features (37). Further, the notion that B cells limit T cell persistence and trafficking has catalyzed a pilot clinical trial of co-infused CD19-targeted chimeric antigen receptor (CAR) T cells with a mesothelin-targeted CAR T cell in PDAC patients (38). Other reports point to presence of B cells with a regulatory phenotype in primary human PDAC, such as those producing IL-10 or IL-35 (39). It will therefore be of interest to delineate whether these B cell features are also evident in metastases as a future objective.

In this study, we also had access to a unique set of matched samples from 4 patients, allowing us to evaluate the direct progression of the PDAC TME and validate trends from the larger cohort. Patients do not typically undergo biopsies of metastasis for routine care, thus even a small cohort such as ours is quite rare and informative, often taking several years to accumulate from pathology archives. While most cells evaluated trended similarly across all patients, the few that diverged may indicate differences in disease process within each patient, or could simply be due to variability because of small sample size. Regardless of their limitations in number, these samples were valuable and informed our data.

Finally, our data show sparse T cell infiltration, particularly of the CD8^+^ T cells in primary PDAC aligns with previous reports (40,41). Strikingly, metastatic tumors had even fewer T cells overall, and that those present were predominantly CD4^+^ lymphocytes. This paucity of T cells suggests that adoptive T cell therapy approaches may be worth pursuing in metastatic PDAC settings.

Transcription factor profiling revealed increased FoxP3 and ROR*γ*t expression in CD4^+^ T cells in primary tumors, consistent with Th2/ Th17 cytokine skewing (42,43), and the presence of Tregs and Th17 cells known to drive PDAC progression (44-46). In contrast, CD4^+^ T cells in metastases expressed more Tbet, indicative of a more inflammatory TME.

We also identified a small population of CD4^+^ CD8^+^ double-positive T cells, previously observed in other cancers but (47,48), to our knowledge, reported here for the first time in PDAC tumors. The concurrent expression of suppressive transcription factors and multiple inhibitory checkpoint receptors on these and other T cells highlights a potential mechanism of resistance to immune checkpoint blockade strategies.

In line with earlier findings in primary PDAC (8), we observed dominant LAG-3, and PD-1 on both CD4^+^ and CD8^+^ T cells. These data support testing rational immunotherapy combinations, such as LAG-3 blockade with myeloid-targeting strategies. Other approaches, such as cancer vaccines, may help mobilize endogenous tumor-specific T cells (49). Ultimately, strategies that improve T cell trafficking, infiltration, and persistence are critical for advancing immunotherapy for PDAC.

This study is valuable to the field as a comprehensive analysis of the TME in primary and metastatic human PDAC, at the protein level. We recognize limitations to this study. We acknowledge the small number of matched samples from patients limits our ability to draw definitive conclusions related to disease process in any one individual. However, as with several other studies utilizing limited patient samples (8,16,20,50), the analysis does uncover unique cellular relationships and is likely to guide future investigation and advances. We hope these data will draw attention to potentially unique roles for myeloid and B cell populations, notably in metastases, that warrant evaluation in larger patient populations. We recognize the limited capacity of mIHC panels to accommodate larger antibody sets, and the focus of our mass cytometry panel on T cell markers prevented obtaining more comprehensive data on other cells in PDAC tumors. For this study, we elected to use CK19 as a well-established marker of malignant epithelial cells, given its routine use in clinical diagnosis of PDAC. Despite these limitations, these results provide valuable insight into specific immune exhaustion patterns of T cells in patient tumors and illuminate redundant mechanisms they may use to circumvent immunotherapy. Our findings provide a rationale for prioritizing emerging targets on tumor-associated macrophages or myeloid-derived suppressor cells in combination with other immunotherapy approaches. We acknowledge that the patients in this study did not receive immunotherapy with either anti-PD-1 or anti-PD-L1 antibodies, consistent with its limited activity in this setting. Future studies including such samples, when appropriate, would provide valuable information regarding the effects of checkpoint blockade therapies on the immune TME and identify changes that may be induced. We also acknowledge that this study is limited to PDAC liver metastasis. While the liver is a common site of metastasis in PDAC, its unique microenvironment is not representative of other metastatic sites and these findings should not be extrapolated to PDAC that has spread to other organs. Future studies could certainly extend our observations to other sites of metastasis or tumor-draining lymph nodes to gather information regarding unique stromal and immune features in a highly aggressive malignancy.

## Ethics Approval

Surgically resected tissue was obtained according to Emory University Institutional Review Board (IRB) protocol IRB00006401 (PI: Krasinskas, last approved: 2/12/2024); and protocol IRB00087397 (PI: Alese, last approved 12/20/2023). The NCT03095781/WCI3321-16 clinical trial was conducted according to Emory University Institutional Review Board (IRB) protocol IRB00087397 (PI: Alese, last approved 12/20/2023). The NCT04191421/WCI4463 clinical trial was conducted according to Emory University Institutional Review Board (IRB) protocol IRB000105616 (PI: Alese, last approved 07/12/2024). Informed consent was obtained from all patients.

## Competing interests

Dr. Lesinski has consulted for ProDa Biotech, LLC, and received compensation. Dr. Lesinski has received research funding through a sponsored research agreement between Emory University and Merck and Co., Bristol-Myers Squibb, Boerhinger-Ingelheim, and Vaccinex. Dr. Paulos has received research funding through a sponsored research agreement between the Medical University of South Carolina and Obsidian, Lycera, ThermoFisher and is the Co-Founder of Ares Immunotherapy. Dr. El-Rayes has reported the following: Research funding from Bristol-Myers Squibb (Inst), Xencor (Inst), Merck Sharp & Dohme (Inst), Exelixis (Inst), GSK (Inst), AstraZeneca (Inst). Consulting or advisory role for AstraZeneca, Genentech. Speaker bureau for Seagen. Dr. Alese has reported the following: Research funding from Taiho Oncology, Ipsen Pharmaceuticals, GSK, Bristol Myers Squibb, SynCore Biotechnology Co. Ltd., Suzhou Transcenta Therapeutics Co., Ltd, Hutchison MediPharma, Boehringer Ingelheim, Xencor Inc., Cue Biopharma, Inc., Merck, Syros Pharmaceuticals Inc., Inhibitex Inc, Arcus Biosciences Inc., ImmunoGen, Impact Therapeutics, Inc. Consulting/Advisory Roles for Ipsen Pharmaceuticals, Aadi Bioscience, Taiho, Pfizer, Seagen Inc., Bristol Myers Squibb, AstraZeneca, Exelixis, Takeda. Independent Data Monitoring Committee for Compass Therapeutics, Inc. Dr. Maithel has reported the following: Funded research from Bristol-Meyers-Squibb and advisory board for AstraZeneca. All other authors declare no competing interests.

## Acknowledgments

We acknowledge the shared resources and cores at Winship Cancer Institute and Emory University that made this research possible including the Winship Cancer Tissue Pathology Shared Resource, the Winship Immune Monitoring Shared Resource, The Winship Data Shared Resource and The Winship Biostatistics Shared Resource *under NIH/NCI award number P30CA138292*. The content is solely the responsibility of the authors and does not necessarily represent the official views of the National Institutes of Health. Figure schematics were created with BioRender.com. We are grateful to Dr. Borgthor Petursson for providing support and computational guidance on spatial analyses and to Dr. Madhav Dhodapkar for valuable discussions surrounding mass cytometry data.

## Funding

This work was supported by NIH grants R01CA228406, R21CA266088-01 (to G. B. Lesinski, B. El-Rayes), and R01CA287866-01 (to G.B. Lesinski, C.M. Paulos), R01CA175061, R01CA208514, R01CA275199 plus Emory University Start-Up Funds (to C.M. Paulos). This work was also supported by the John Kauffman Family Professorship for Pancreatic Cancer Research (to G.B. Lesinski). NKH, PhD is supported by a Postdoctoral Fellowship from the American Cancer Society, https://doi.org/10.53354/ACS.PF-24-1258662-01-IBCD.pc.gr.222223.

## Author Contributions

Conceptualization: EG, NKH, HTK, BFE, ATR, CMP, and GBL; Sample Acquisition: AMK, SKM, JMS, MMS, MYZ, MD, BFE, OBA; Clinical Trial Oversight: MD, OBA, BFE; Methodology: EG, NKH, DBD, VCP, JK, CJH, EEG, and ATR; Investigation: EG, DBD, VCP, and NKH; Formal analysis and interpretation of data: EG, NKH, ATR, KD, HTK, CMP and GBL; Resources: BE, CMP and GBL; Consultation: ATK, KD, HTK and CMP; Visualization: EG and NKH; Funding acquisition: BFE, CMP, GBL; Supervision: GBL; Guarantor of the project: GBL. All authors performed writing (review and editing) and agreed on its content.

