## Supplemental Material for "High-Dimensional Protein Analysis Uncovers Distinct Immunological and Stromal Signatures Between Primary and Metastatic Pancreatic Ductal Adenocarcinoma"

### Figure S1

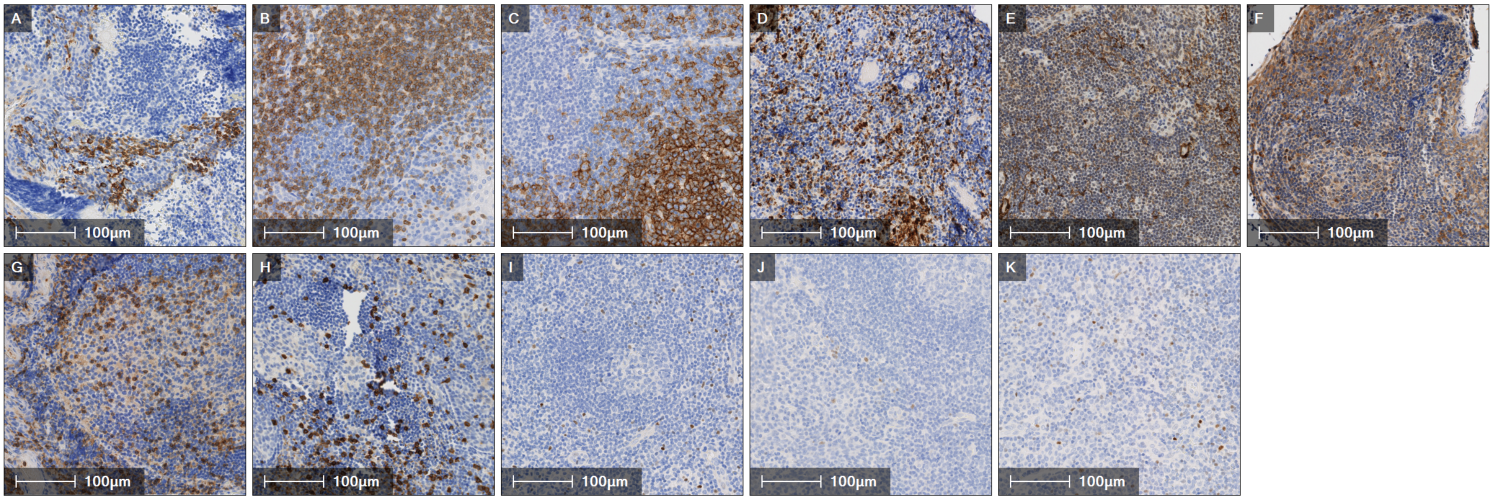
A
100µm
B
100µm
C
100µm
D
100µm
E
100µm
F
100µm
G
100µm
H
100µm
I
100µm
J
100µm
K
100µm

**Supplemental Figure 1** Single stain IHC used for validation of multiplex IHC panels. Human tonsil tissue was used as a biologic control for validating single antibody staining prior to use on multiplex IHC staining panels. Representative images are provided for each stain and are arranged in the figure as follows: (A) CK19, (B) CD3, (C) CD19, (D) CD68, (E) α-SMA, (F) IL-6, (G) CD4, (H) CD8, (I) FoxP3, (J) RORγt and (K) T-bet.

### Figure S2

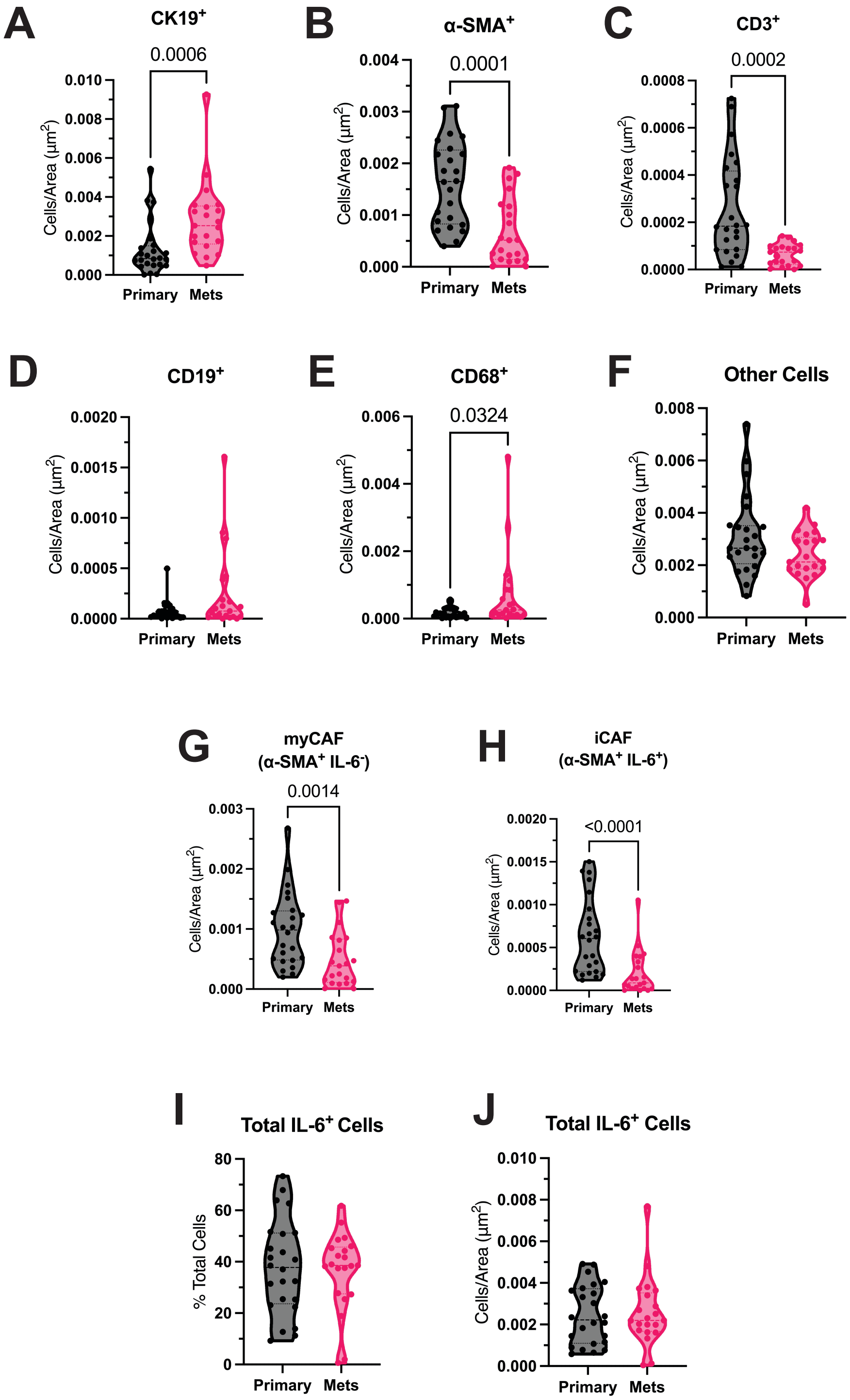
A
CK19+
0.0006
Cells/Area (µm2)
Primary
Mets
B
α-SMA+
0.0001
C
CD3+
0.0002
D
CD19+
E
CD68+
0.0324
F
Other Cells
G
myCAF
(α-SMA+ IL-6-)
0.0014
H
iCAF
(α-SMA+ IL-6+)
<0.0001
I
Total IL-6+ Cells
% Total Cells
J
Total IL-6+ Cells

**Supplemental Figure 2** Density of phenotyped cells in tissue based on mIHC analysis validates total cell quantification. Truncated violin plots show the abundance of the following cell types: (A) $CK19^+$, (B) α-$SMA^+$, (C) $CD3^+$, (D) $CD19^+$, (E) $CD68^+$, (F) other $DAPI^+$, (G) myCAF (α-$SMA^+$), and (H) iCAF (α-$SMA^+$IL-$6^+$) in primary and metastatic tissue, relative to tissue area ($μm^2$). Truncated violin plots show the abundance of total IL-$6^+$ cells as (I) a percent of total cells and (J) density ($μm^2$) in primary and metastatic tissue. Statistical comparisons were done using Mann-Whitney tests. Sample size: n=24, primary; n=21, metastatic.

### Figure S3

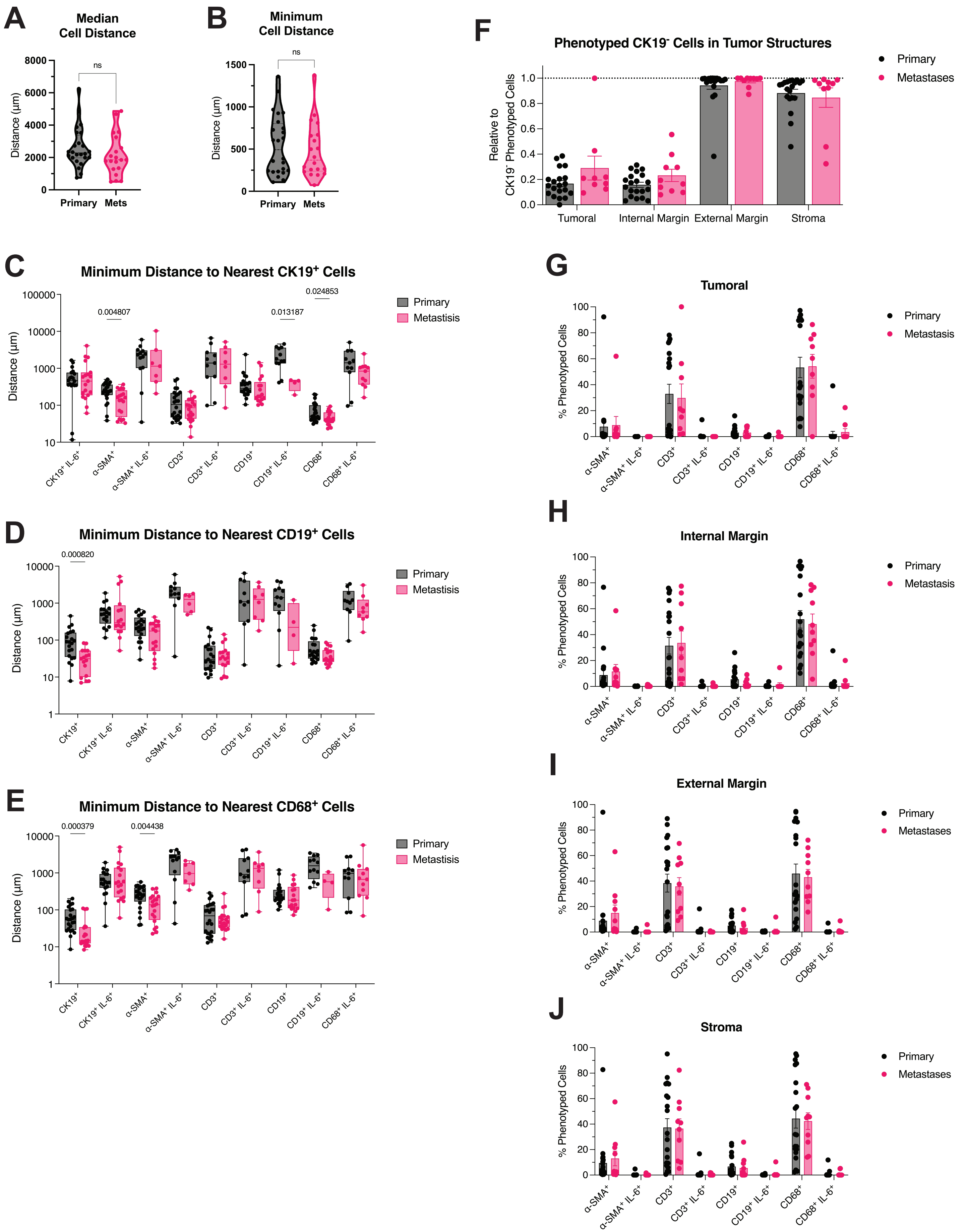

A
Median Cell Distance
ns
Distance (µm)
8000
6000
4000
2000
0
Primary
Mets
B
Minimum Cell Distance
ns
Distance (µm)
1500
1000
500
0
Primary
Mets
F
Phenotyped CK19⁻ Cells in Tumor Structures
Relative to CK19⁻ Phenotyped Cells
1.0
0.8
0.6
0.4
0.2
0.0
Tumoral
Internal Margin
External Margin
Stroma
Primary
Metastases
C
Minimum Distance to Nearest CK19⁺ Cells
Distance (µm)
100000
10000
1000
100
10
0.004807
0.013187
0.024853
CK19⁺ IL-6⁺
α-SMA⁺
α-SMA⁺ IL-6⁺
CD3⁺
CD3⁺ IL-6⁺
CD19⁺
CD19⁺ IL-6⁺
CD68⁺
CD68⁺ IL-6⁺
Primary
Metastisis
D
Minimum Distance to Nearest CD19⁺ Cells
Distance (µm)
10000
1000
100
10
1
0.000820
CK19⁺
CK19⁺ IL-6⁺
α-SMA⁺
α-SMA⁺ IL-6⁺
CD3⁺
CD3⁺ IL-6⁺
CD19⁺ IL-6⁺
CD68⁺
CD68⁺ IL-6⁺
Primary
Metastisis
E
Minimum Distance to Nearest CD68⁺ Cells
Distance (µm)
10000
1000
100
10
1
0.000379
0.004438
CK19⁺
CK19⁺ IL-6⁺
α-SMA⁺
α-SMA⁺ IL-6⁺
CD3⁺
CD3⁺ IL-6⁺
CD19⁺
CD19⁺ IL-6⁺
CD68⁺ IL-6⁺
Primary
Metastisis
G
Tumoral
% Phenotyped Cells
100
80
60
40
20
0
α-SMA⁺
α-SMA⁺ IL-6⁺
CD3⁺
CD3⁺ IL-6⁺
CD19⁺
CD19⁺ IL-6⁺
CD68⁺
CD68⁺ IL-6⁺
Primary
Metastasis
H
Internal Margin
% Phenotyped Cells
100
80
60
40
20
0
α-SMA⁺
α-SMA⁺ IL-6⁺
CD3⁺
CD3⁺ IL-6⁺
CD19⁺
CD19⁺ IL-6⁺
CD68⁺
CD68⁺ IL-6⁺
Primary
Metastasis
I
External Margin
% Phenotyped Cells
100
80
60
40
20
0
α-SMA⁺
α-SMA⁺ IL-6⁺
CD3⁺
CD3⁺ IL-6⁺
CD19⁺
CD19⁺ IL-6⁺
CD68⁺
CD68⁺ IL-6⁺
Primary
Metastases
J
Stroma
% Phenotyped Cells
100
80
60
40
20
0
α-SMA⁺
α-SMA⁺ IL-6⁺
CD3⁺
CD3⁺ IL-6⁺
CD19⁺
CD19⁺ IL-6⁺
CD68⁺
CD68⁺ IL-6⁺
Primary
Metastases

**Supplemental Figure 3** Spatial analysis reveals distinct cellular distribution and proximity of immune cells to $CK19^+$ cells between primary and metastatic PDAC tumors. (A) Truncated violin plot quantifying median distances between cells, providing overall cell spacing across each tissue sample (Mann-Whitney test). (B) Truncated violin plot quantifying minimum distances between cells, providing insights into cell-to-cell interactions, where proximity is critical for function or signaling (Mann-Whitney test). Box and whisker plots quantifying minimum distance of individual cell types to (C) $CK19^+$, (D) $CD19^+$ and (E) $CD68^+$ cells within primary and metastatic PDAC tumors (Mann-Whitney tests). (F) Bar graph quantifying the relative fraction of $CK19^-$ cells in the four defined tumor structures. (G-J) Bar graph quantifying percent of each cell type in the tumoral, internal margin, external margin, and stroma regions (Mann-Whitney test). Sample sizes: tissue spatial analysis: n=24, primary; n=20, metastatic; tumor structure analysis: n=20, primary; n=10, metastatic.

### Figure S4

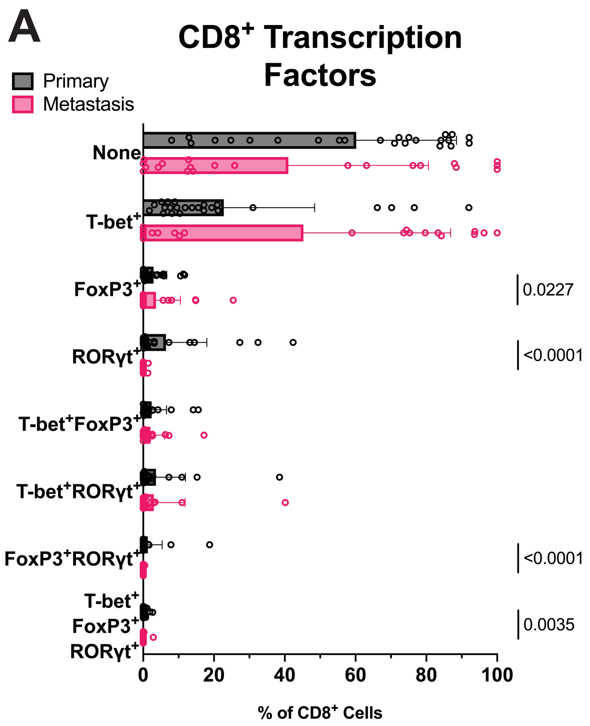

A
CD8+ Transcription Factors
Primary
Metastasis
None
T-bet+
FoxP3+
RORγt+
T-bet+FoxP3+
T-bet+RORγt+
FoxP3+RORγt+
T-bet+ FoxP3+ RORγt+
0.0227
<0.0001
<0.0001
0.0035
0
20
40
60
80
100
% of CD8+ Cells

**Supplemental Figure 4** Transcription factor levels in $CD8^+$ T cells in tissues quantified from mIHC images reveals a trend towards suppressive phenotypes. (A) Bar graph showing the mean and standard deviation (SD) of the percentage of transcription factor (T-bet, FoxP3 and ROR$\gamma$t) positive $CD8^+$ T cell populations represented as a percent of total $CD8^+$ T cells (Mann-Whitney tests). Sample sizes: n=24, primary; n=21, metastatic.

### Figure S5

A

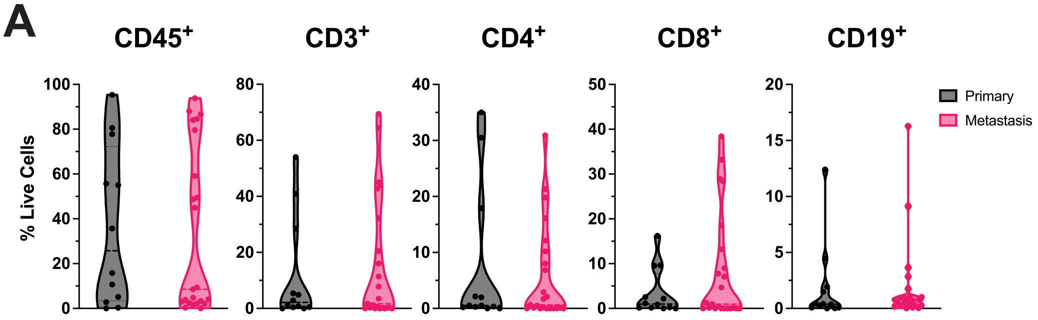

CD45+
CD3+
CD4+
CD8+
CD19+
% Live Cells
100
80
60
40
20
0
Primary
Metastasis


B

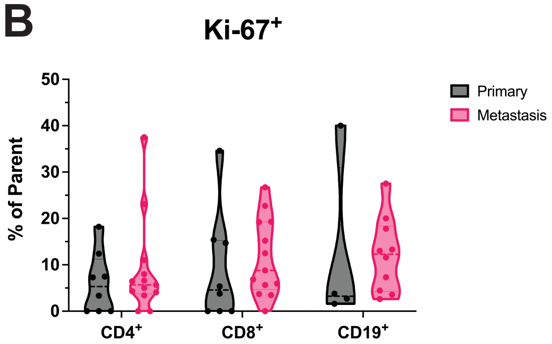

Ki-67+
% of Parent
50
40
30
20
10
0
CD4+
CD8+
CD19+
Primary
Metastasis

**Supplemental Figure 5** Quantification for select lymphocyte populations by mass cytometry indicates greater proliferation in metastatic tissue. (A) Truncated violin plots of lymphocyte populations detected shown as a percentage of total live cells (Mann-Whitney test). Sample size: n=12, primary; n=23, metastatic. (B) Truncated violin plots quantifying the expression of Ki-67 on $CD4^+$, $CD8^+$ and $CD19^+$ cells as a percent of total $CD4^+$, $CD8^+$ or $CD19^+$ cells, respectively. $CD4^+$ and $CD8^+$ sample size: n=8, primary; n=13, metastatic. $CD19^+$ sample size: n=4, primary; n=10, metastatic.

### Figure S6

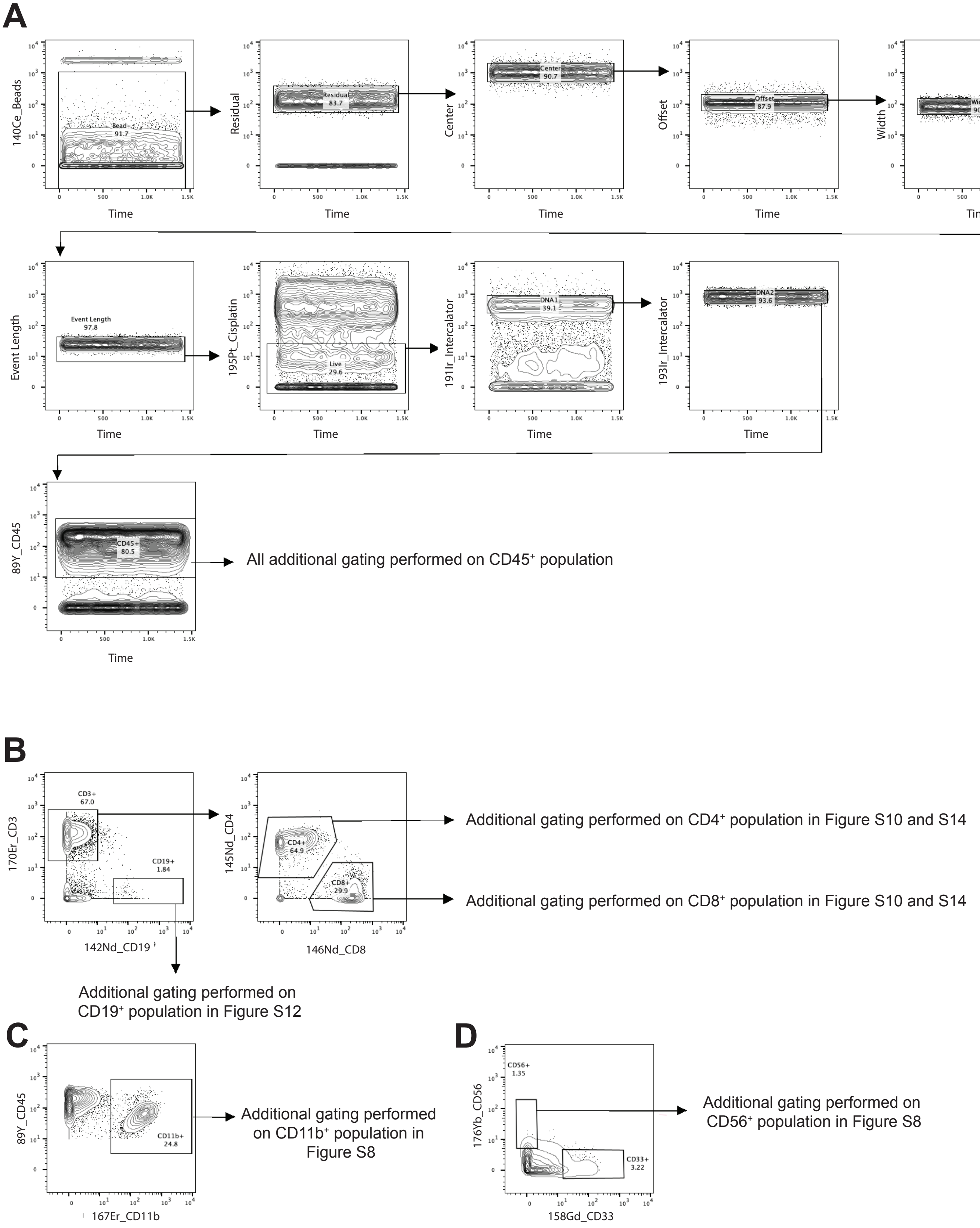
A
140Ce_Beads
Bead
91.7
Time
Residual
83.7
Center
90.7
Offset
87.9
Width
90.9
Event Length
97.8
195Pt_Cisplatin
Live
29.6
191Ir_Intercalator
DNA1
39.1
193Ir_Intercalator
DNA2
93.6
89Y_CD45
CD45+
80.5
All additional gating performed on CD45⁺ population
B
170Er_CD3
CD3+
67.0
CD19+
1.84
142Nd_CD19
145Nd_CD4
CD4+
64.9
CD8+
29.9
146Nd_CD8
Additional gating performed on CD4⁺ population in Figure S10 and S14
Additional gating performed on CD8⁺ population in Figure S10 and S14
Additional gating performed on CD19⁺ population in Figure S12
C
89Y_CD45
CD11b+
24.8
167Er_CD11b
Additional gating performed on CD11b⁺ population in Figure S8
D
176Yb_CD56
CD56+
1.35
CD33+
3.22
158Gd_CD33
Additional gating performed on CD56⁺ population in Figure S8

**Supplemental Figure 6** Representative gating scheme for the analysis of mass cytometry data used throughout the manuscript. (A) $CD45^+$ cells were gated based on live single cell population. (B) Identification of $CD3^+$ and $CD19^+$ cells from $CD45^+$ cells, followed by selection of $CD4^+$ and $CD8^+$ cells out of $CD3^+$ parent population. (C) Identification of $CD11b^+$ cells from $CD45^+$ cells. (D) Identification of $CD56^+$ out of $CD45^+$ cells. Representative sample: P7.

### Figure S7

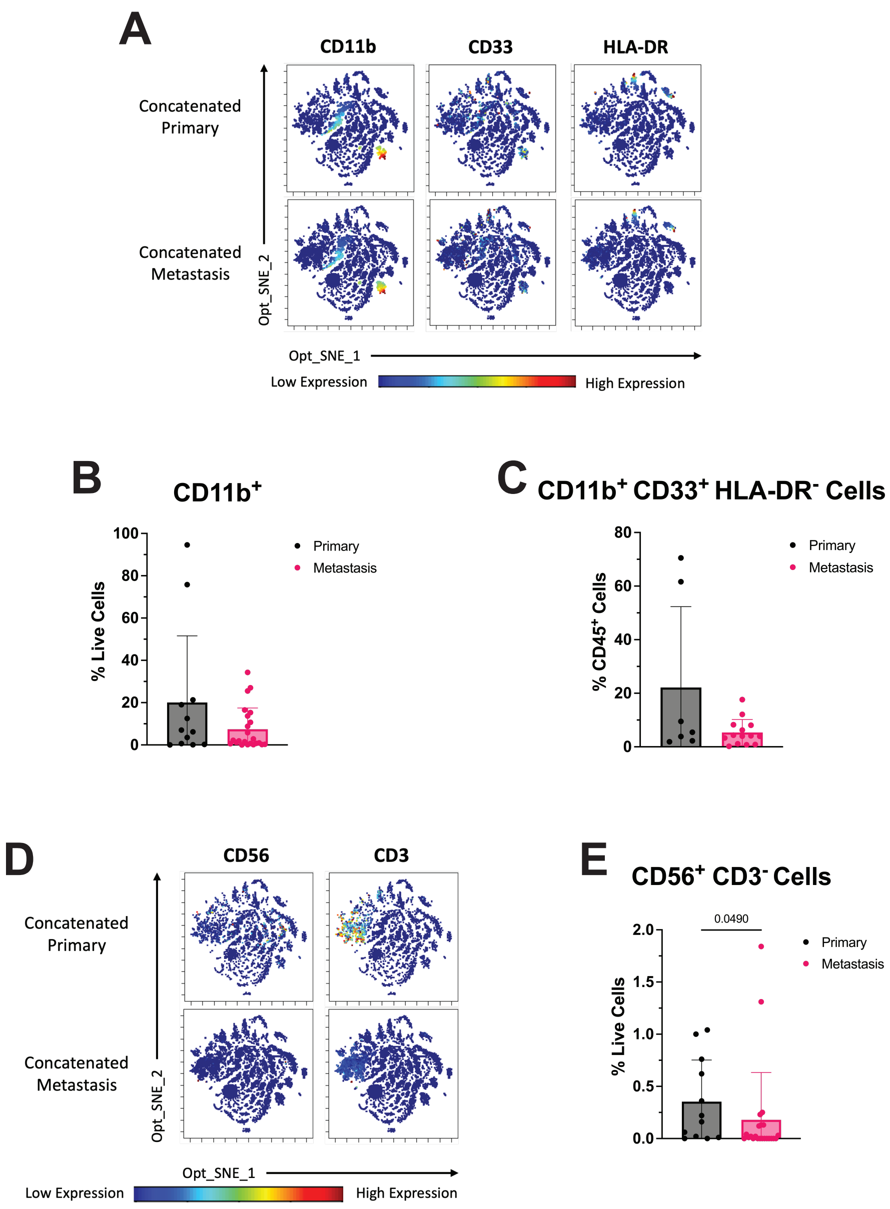
A
CD11b
CD33
HLA-DR
Concatenated Primary
Concatenated Metastasis
Opt_SNE_2
Opt_SNE_1
Low Expression
High Expression
B
CD11b+
% Live Cells
100
80
60
40
20
0
Primary
Metastasis
C
CD11b+ CD33+ HLA-DR- Cells
% CD45+ Cells
80
60
40
20
0
Primary
Metastasis
D
CD56
CD3
Concatenated Primary
Concatenated Metastasis
Opt_SNE_2
Opt_SNE_1
Low Expression
High Expression
E
CD56+ CD3- Cells
0.0490
% Live Cells
2.0
1.5
1.0
0.5
0.0
Primary
Metastasis

**Supplemental Figure 7** Phenotypically defined myeloid and natural killer cells using mass cytometry reveals differences in cellular composition between primary and metastatic PDAC. (A) Levels of CD11b, CD33, and HLA-DR displayed on individual, unsupervised Opt_SNE plots of live single cells in concatenated primary (top) and metastatic (bottom) samples. Bar graph with mean and SD quantification of (B) $CD11b^+$ cells as a percent of live cells and (C) myeloid cells ($CD11b^+$ $CD33^+$ $HLA\text{-}DR^+$) as a percent of $CD45^+$ cells. (D) Expression of CD56 and CD3 shown in individual, unsupervised Opt_SNE plots of live single cells in concatenated primary (top) and metastatic (bottom) samples. (E) Bar graph with mean and SD quantification of natural killer cells ($CD56^+$ $CD3^-$) (Mann-Whitney test). Myeloid cell sample sizes: n=7, primary; n=14, metastatic. $CD11b^+$ and $CD56^+CD3^-$ cell sample size: n=12, primary; n=23, metastatic.

### Figure S8

**A**

Continued from Figure S6C

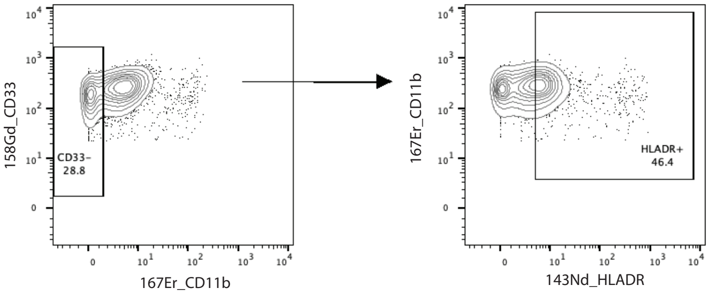


Continued from Figure S6D

**B**

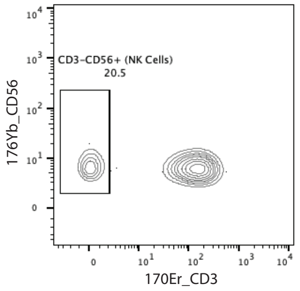

**Supplemental Figure 8** Mass cytometry gating strategy of myeloid and natural killer cell populations. Representative plots show subpopulations identified by (A) CD11b$^+$, CD33$^+$, HLA-DR$^+$ cells and (B) CD3$^-$ CD56$^+$ cells. Representative sample: P7.

### Figure S9

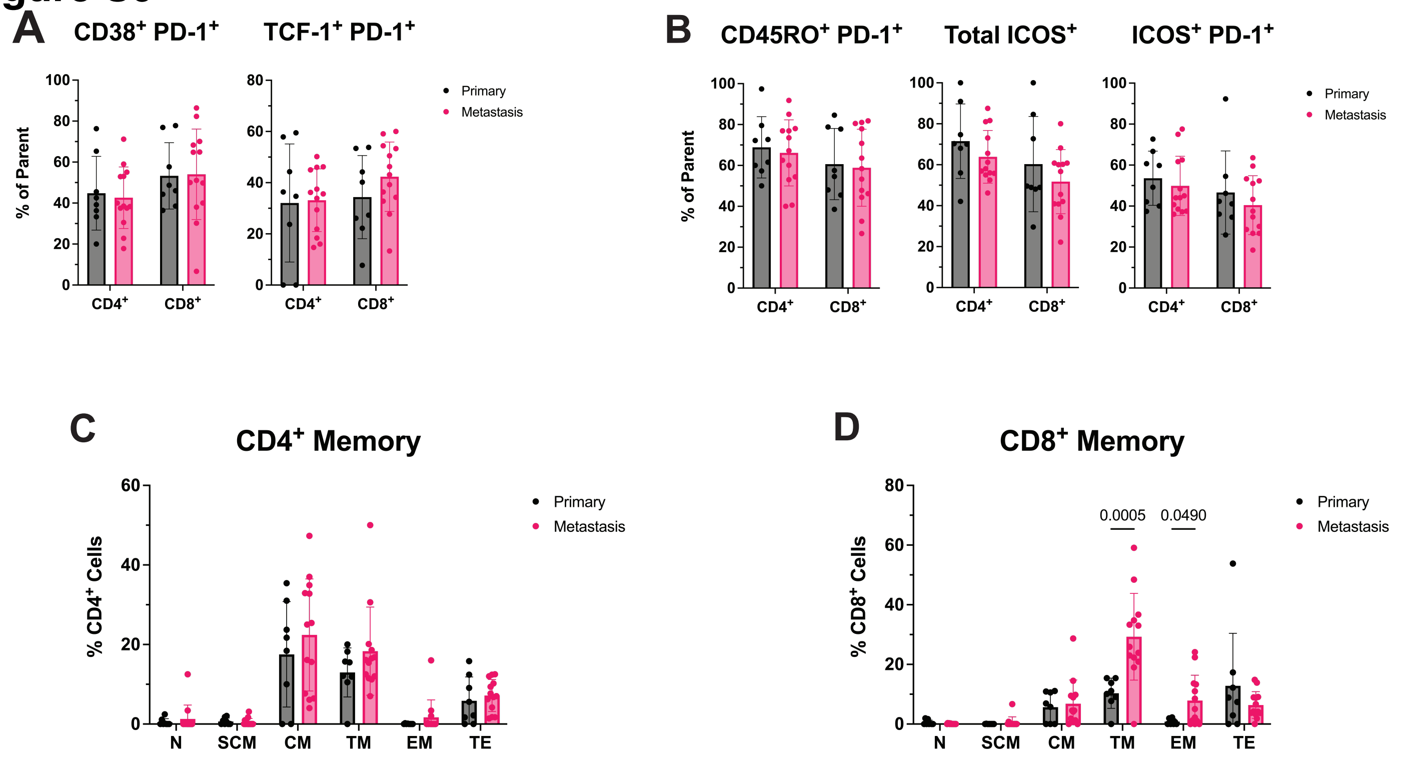
A
CD38+ PD-1+
TCF-1+ PD-1+
% of Parent
CD4+
CD8+
Primary
Metastasis
B
CD45RO+ PD-1+
Total ICOS+
ICOS+ PD-1+
% of Parent
CD4+
CD8+
Primary
Metastasis
C
CD4+ Memory
% CD4+ Cells
N
SCM
CM
TM
EM
TE
Primary
Metastasis
D
CD8+ Memory
% CD8+ Cells
0.0005
0.0490
N
SCM
CM
TM
EM
TE
Primary
Metastasis

**Supplemental Figure 9** T cell populations in metastatic PDAC suggest greater antigen exposure and exhaustion than primary tumors. Bar graphs with mean and SD quantification subsets of $CD4^+$ and $CD8^+$ T cells expressing (A) $CD38^+$ or TCF-$1^+$ with PD-$1^+$ and (B) $CD45RO^+$ PD-$1^+$, $ICOS^+$ with and without PD-$1^+$, as well as, memory T cell subsets (N, SCM, CM, TM, EM and TE) for (C) $CD4^+$ and (D) $CD8^+$ T cells, respectively. $CD4^+$ and $CD8^+$ T cell sample size: n=8, primary; n=13, metastatic. N, naïve ($CD45RO^-$ $CD127^+$ $CD27^+$ $CCR7^+$ $CD95^-$); SCM, stem-like central memory ($CD45RO^-$ $CD127^+$ $CD27^+$ $CCR7^+$ $CD95^+$); CM, central memory ($CD45RO^+$ $CD127^+$ $CD27^+$ $CCR7^+$ $CD95^+$); TM, transitional memory ($CD45RO^+$ $CCR7^-$ $CD27^+$ $CD95^+$); EM, effector memory ($CD45RO^-$ $CD127^-$ $CD27^-$ $CD95^+$); TE, terminal effector ($CD45RO^+$ $CCR7^-$ $CD27^-$ $CD95^+$ $CD127^-$).

### Figure S10

Continued from Figure S6B

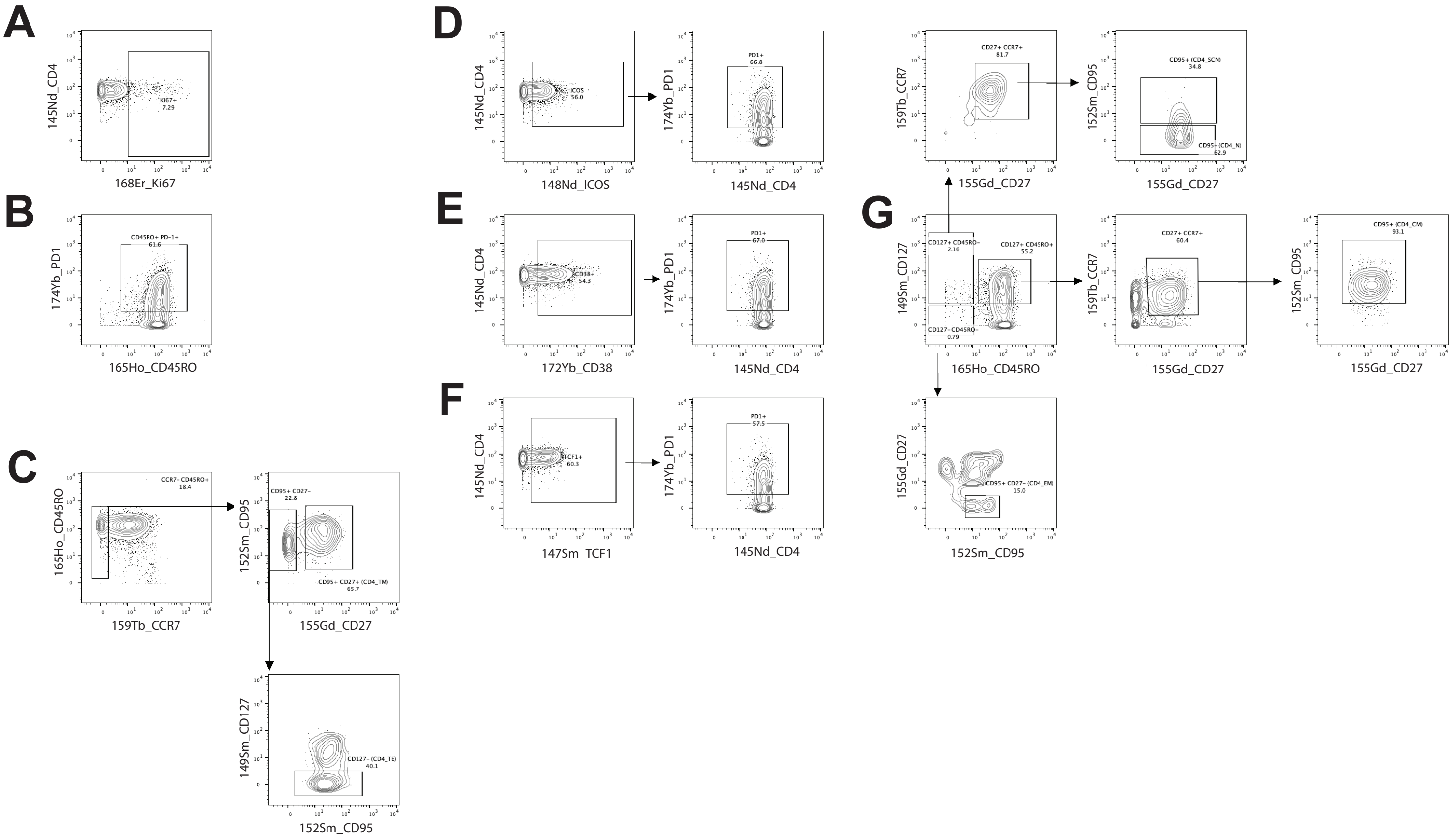


Continued from Figure S6B

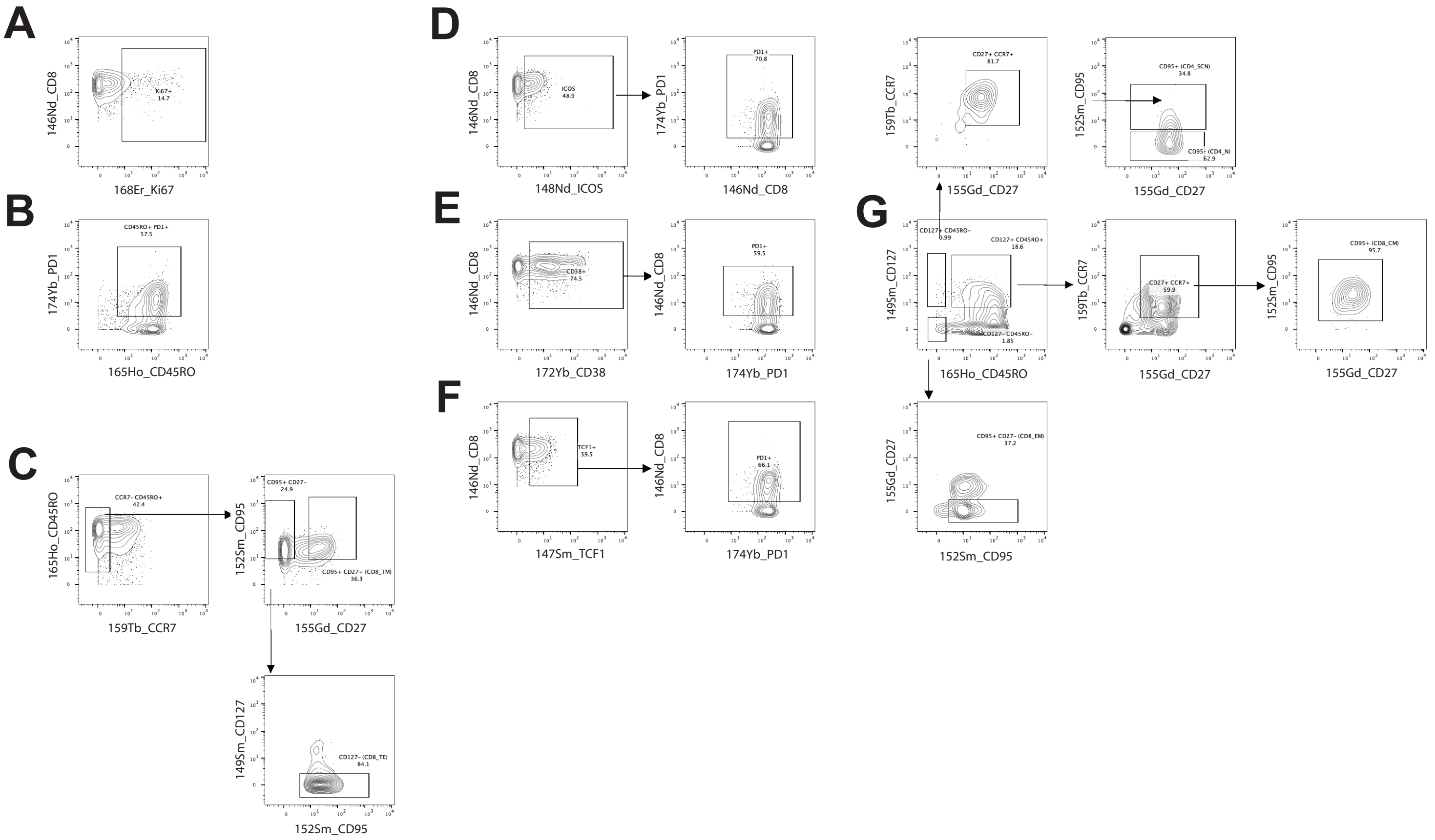

**Supplemental Figure 10** Mass cytometry gating strategy of $CD4^+$ and $CD8^+$ T cell subpopulations. Subpopulations include (A) Ki-$67^+$, (B) $CD45RO^+$ PD-$1^+$, (C) $CCR7^-$ $CD45RO^+$ $CD95^+$ $CD27^-$ $CD127^-$, TE cells and $CCR7^-$ $CD45RO^+$ $CD95^+$ $CD27^+$, TM cells, (D) $ICOS^+$ PD-$1^+$, (E) $CD38^+$ PD-$1^+$, (F) TCF-$1^+$ PD-$1^+$, (G) $CD127^+$ $CD45RO^-$ $CD27^+$ $CCR7^+$ $CD95^+$, SCN cells; $CD127^+$ $CD45RO^-$ $CD27^+$ $CCR7^+$ $CD95^-$, N cells; $CD127^+$ $CD45RO^+$ $CD27^+$ $CCR7^+$ $CD95^+$, CM cells; $CD127^-$ $CD45RO^-$ $CD27^-$ $CD95^+$, EM cells. Representative sample: P7. TE, terminal effector; TM, transitional memory; SCM, stem-like central memory; N, naïve; CM, central memory; EM, effector memory.

### Figure S11

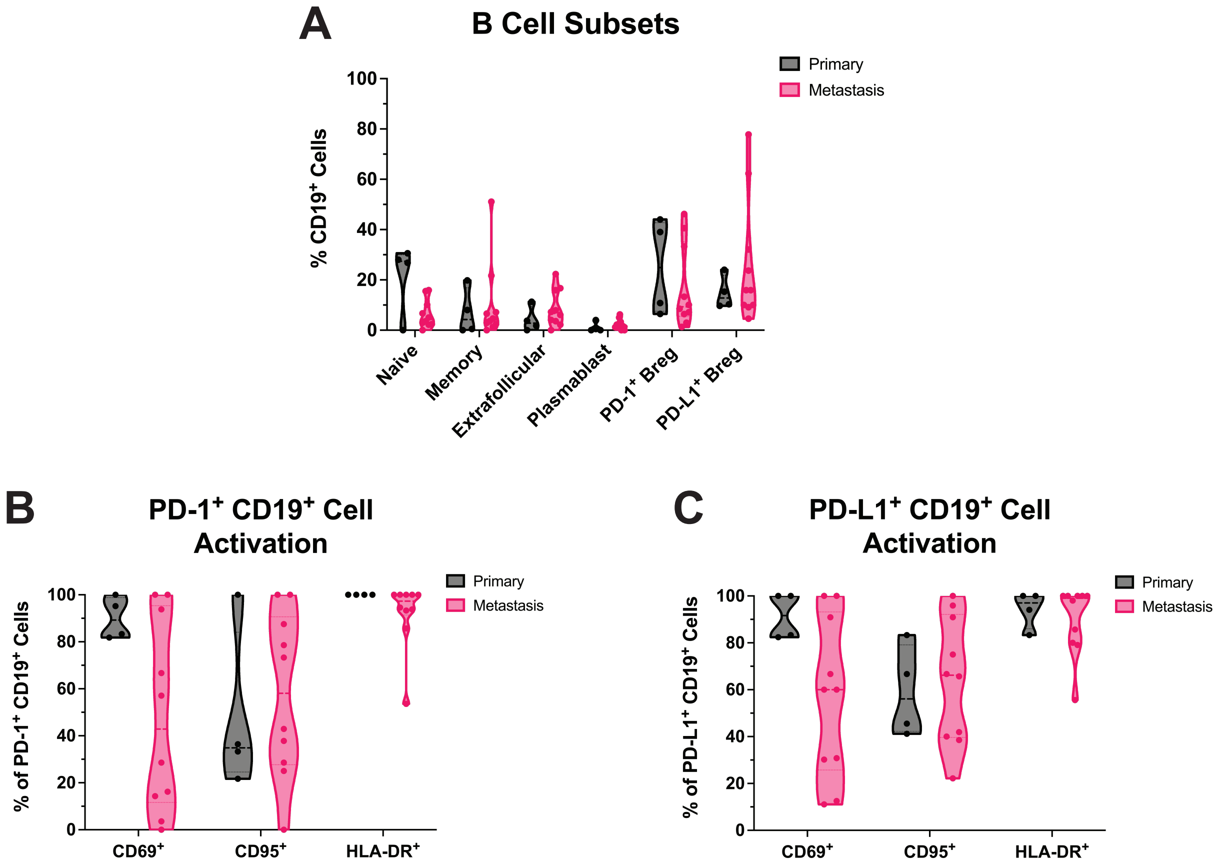
A
B Cell Subsets
Primary
Metastasis
% CD19+ Cells
100
80
60
40
20
0
Naive
Memory
Extrafollicular
Plasmablast
PD-1+ Breg
PD-L1+ Breg
B
PD-1+ CD19+ Cell Activation
% of PD-1+ CD19+ Cells
CD69+
CD95+
HLA-DR+
C
PD-L1+ CD19+ Cell Activation
% of PD-L1+ CD19+ Cells
CD69+
CD95+
HLA-DR+

**Supplemental Figure 11** B cell subsets and their activation markers within PDAC. (A) Truncated violin plots of naïve, $CD27^+$ Memory, Extrafollicular, Plasmablast, $PD\text{-}1^+$ and $PD\text{-}L1^+$ B cell subsets shown as a percentage of $CD19^+$ population in tissue samples. Truncated violin plots quantify $CD69^+$, $CD95^+$ and $HLA\text{-}DR^+$ of (B) $PD\text{-}1^+$ $CD19^+$ and (C) $PD\text{-}L1^+$ $CD19^+$ cells, represented as a percent of parent cell. $CD19^+$ sample size: n=4, primary; n=10, metastatic.

### Figure S12

Continued from Figure S6B

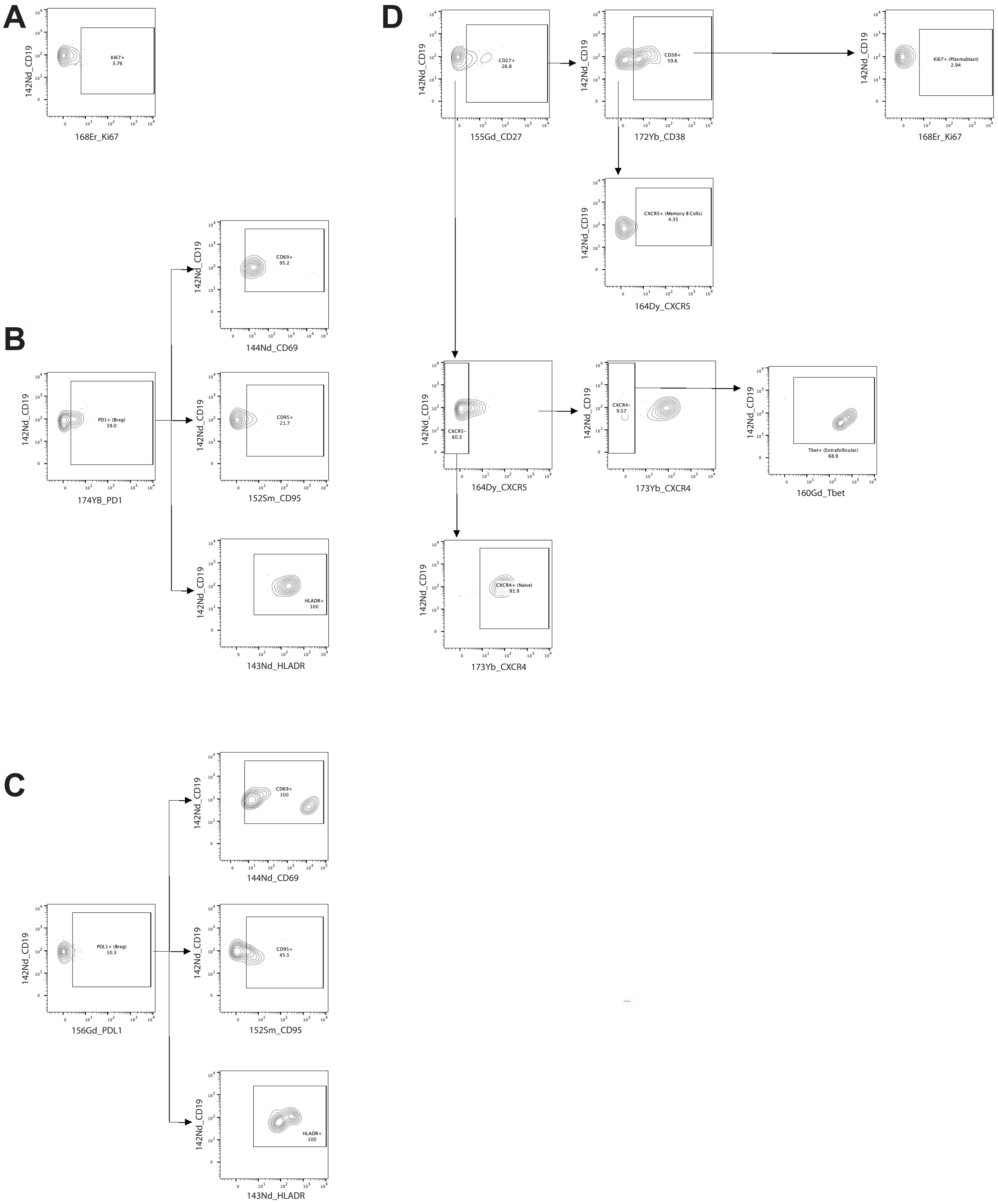

**Supplemental Figure 12** Mass cytometry gating strategy of $CD19^+$ cell subpopulations. Representative plots of identified subpopulations including (A) Ki-$67^+$, (B) PD-$1^+$ Breg, $CD69^+$, $CD95^+$ and HLA-$DR^+$ cells, (C) PD-$L1^+$ Breg, $CD69^+$, $CD95^+$ and HLA-$DR^+$ cells, and (D) $CD27^+$, $CD38^+$, Ki-$67^+$, Plasmablasts; $CD27^+$, $CD38^-$, $CXCR5^+$, Memory B cells; $CD27^-$, $CXCR5^+$, $CXCR4^-$, T-$bet^+$, Extrafollicular B cells; $CD27^-$, $CXCR5^-$, $CXCR4^+$, Naïve B cells. Representative sample: P7. Bregs, B regulatory cells.

### Figure S13

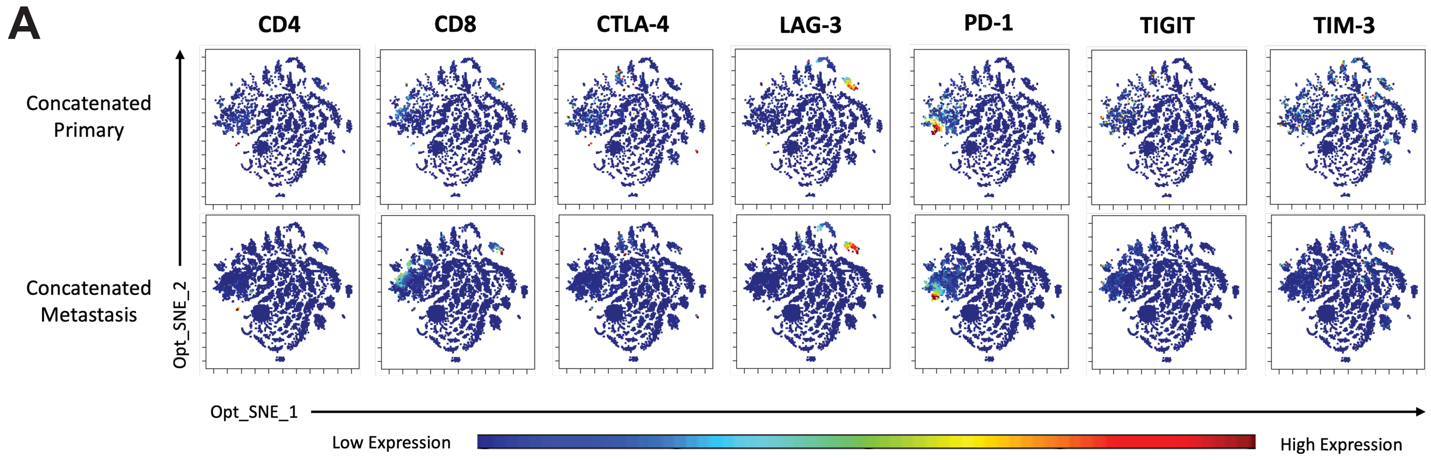
A
CD4
CD8
CTLA-4
LAG-3
PD-1
TIGIT
TIM-3
Concatenated Primary
Concatenated Metastasis
Opt_SNE_2
Opt_SNE_1
Low Expression
High Expression

**Supplemental Figure 13** Opt_SNE plots reveal abundant checkpoint marker expression on $CD4^+$ and $CD8^+$ T cells. (A) Expression of CTLA-4, LAG-3, PD-1, TIGIT and TIM-3, along with CD4 and CD8, displayed on individual, unsupervised Opt_SNE plots of live single cells in concatenated primary (top) and metastatic (bottom) samples. Opt_SNE, optimizes t-distributed stochastic neighbor embedding.

### Figure S14

Continued from Figure S6B

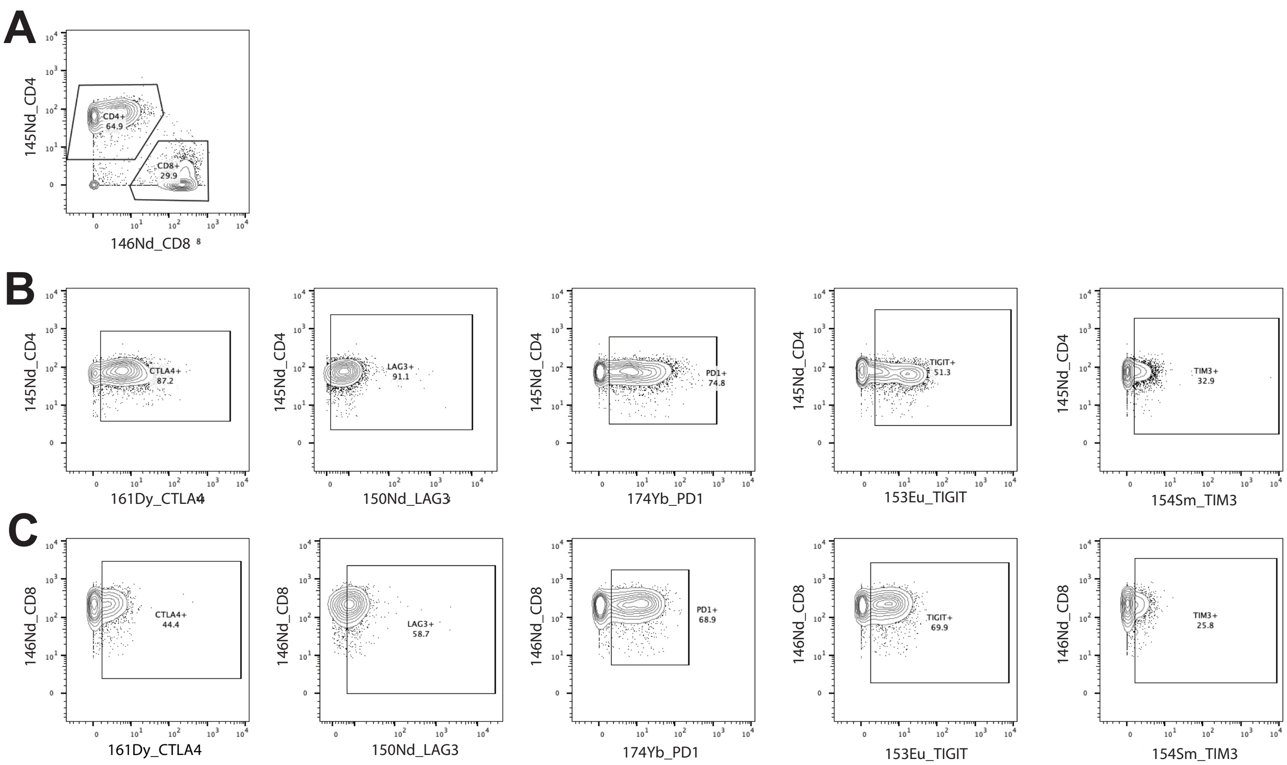

**Supplemental Figure 14** Mass cytometry gating strategy of inhibitory immune checkpoint receptors present on $CD4^+$ and $CD8^+$ T cell populations. (A) $CD4^+$ and $CD8^+$ T cells selected from $CD3^+$ population. Representative plots of the checkpoint molecules CTLA-4, LAG-3, PD-1, TIGIT, and TIM-3 on (B) $CD4^+$ cells and (C) $CD8^+$ cells, respectively. Representative sample: P8.

### Figure S15

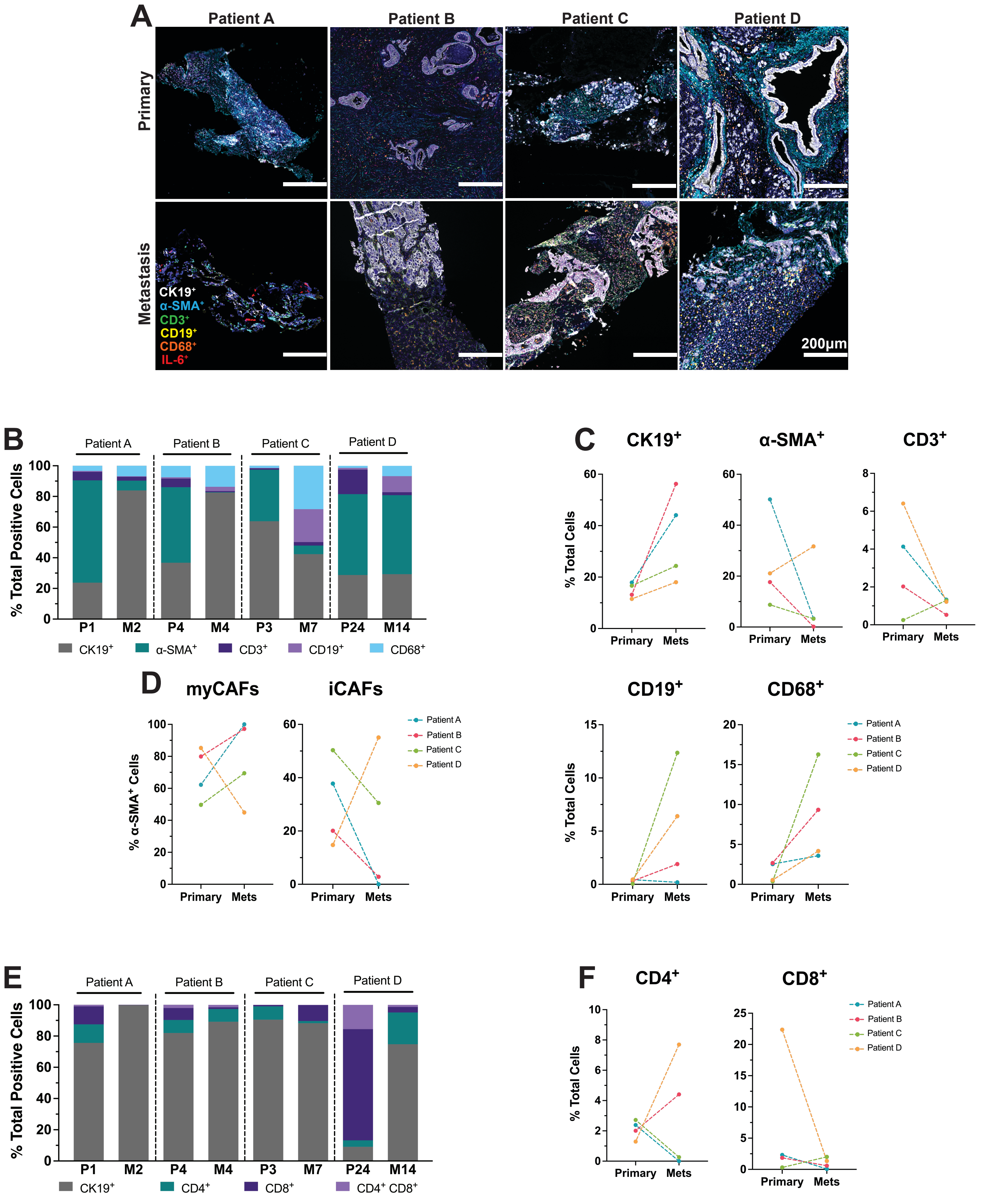

A
Patient A
Patient B
Patient C
Patient D
Primary
Metastasis
CK19+
α-SMA+
CD3+
CD19+
CD68+
IL-6+
200μm
B
% Total Positive Cells
P1 M2 P4 M4 P3 M7 P24 M14
CK19+ α-SMA+ CD3+ CD19+ CD68+
C
CK19+
α-SMA+
CD3+
CD19+
CD68+
% Total Cells
Primary Mets
D
myCAFs
iCAFs
% α-SMA+ Cells
Patient A
Patient B
Patient C
Patient D
E
% Total Positive Cells
P1 M2 P4 M4 P3 M7 P24 M14
CK19+ CD4+ CD8+ CD4+ CD8+
F
CD4+
CD8+
% Total Cells
Primary Mets

**Supplemental Figure 15** Matched primary and metastatic PDAC samples validates trends of larger cohort analysis. (A) Representative images of mIHC staining on three primary (top) and patient-matched metastatic (bottom) tumor samples. Antibody panel detecting IL-6 (pink), CD68 (red), α-SMA (cyan), CK19 (white), CD19 (yellow) and CD3 (green) fluorescent signals. (B) Stacked bar graph showing the distribution of $CK19^+$, $α\text{-}SMA^+$, $CD3^+$, $CD19^+$, and $CD68^+$ cells detected by mIHC in each patient sample. Lines plots comparing (C) $CK19^+$, $α\text{-}SMA^+$, $CD3^+$, $CD19^+$, and $CD68^+$ or (D) myCAF and iCAF subtypes in matched primary and metastatic tumors of each patient. (E) Stacked bar graph of showing the distribution of $CK19^+$, $CD4^+$, $CD8^+$ and $CD4^+$ $CD8^+$ cells detected by mIHC in each patient sample. Before-after plots with lines representing each patient quantifying (F) $CD4^+$ and $CD8^+$ T cells. n=4 primary and metastatic matched samples.

| **Supplemental Table 1: Patient Demographics** | |
|---|---|
| **Primary Resections** | **N=27** |
| **Age** | |
| >=48, <65 | 14 |
| >=65, <79 | 13 |
| **Sex** | |
| Female | 15 |
| Male | 12 |
| **Race** | |
| Black | 7 |
| White | 17 |
| Other | 2 |
| Unknown | 1 |
| **Prior Therapies** | |
| 1 | 15 |
| 2 | 7 |
| >=3 | 5 |
| **Tumor Site** | |
| Ampullary | 1 |
| Uncinate Process | 1 |
| Pancreatic Head | 17 |
| Pancreatic Body | 4 |
| Pancreatic Tail | 2 |
| Not Specified | 2 |

| **Supplemental Table 2: Patient Demographics** | |
|---|---|
| **Metastatic Liver Biopsies** | **N=26** |
| **Age** | |
| >=48, <65 | 12 |
| >=65, <81 | 14 |
| **Sex** | |
| Female | 11 |
| Male | 15 |
| **Race** | |
| Black | 4 |
| White | 18 |
| Other | 1 |
| Unknown | 3 |
| **Prior Therapies** | |
| 1 | 2 |
| 2 | 12 |
| >=3 | 12 |

**Supplemental Table 3. Primary Sample Cell Count**

| Sample | mIHC | | | CyTOF |
|---|---|---|---|---|
| | Cell Detections | Tissue area (µm^2) | Cell/Area Density (cells per µm^2) | Live Single Cells |
| P1 | 22994 | 6955331 | 302 | - |
| P2 | 2572850 | 217565314 | 85 | - |
| P3 | 160246 | 35178257 | 220 | - |
| P4 | 3024033 | 325113832 | 108 | - |
| P5 | 2739416 | 236145193 | 86 | - |
| P6 | 1212971 | 129313306 | 107 | - |
| P7 | 301728 | 44194497 | 146 | 14380 |
| P8 | 190350 | 28655434 | 151 | 4658 |
| P9 | 141223 | 19095375 | 135 | 5388 |
| P10 | 93104 | 12181366 | 131 | 3076 |
| P11 | - | - | - | 2493 |
| P12 | - | - | - | 114 |
| P13 | 56737 | 9866383 | 174 | - |
| P14 | 72170 | 13183450 | 183 | - |
| P15 | 146119 | 43451128 | 297 | - |
| P16 | 133154 | 52780782 | 396 | - |
| P17 | 174896 | 27069969 | 155 | - |
| P18 | 148521 | 28976871 | 195 | - |
| P19 | 98657 | 18401399 | 187 | 135 |
| P20 | 351011 | 43702588 | 125 | 514 |
| P21 | 84327 | 12614542 | 150 | 1264 |
| P22 | 120054 | 18681141 | 156 | 502 |
| P23 | 99269 | 13467538 | 136 | 1156 |
| P24 | 952499 | 134809771 | 142 | - |
| P25 | 255852 | 67792533 | 265 | 79779 |
| P26 | - | - | - | 34971 |
| P27 | 6286 | 1270986 | 202 | 17017 |

*P=Primary

**Supplemental Table 4. Metastasis Sample Cell Count**

| Sample | mIHC | | | CyTOF |
|---|---|---|---|---|
| | Cell Detections | Tissue area (μm^2) | Cell/Area Density (cells per μm^2) | Live Single Cells |
| M1 | - | - | - | 695 |
| M2 | 2178 | 291250 | 134 | 2088 |
| M3 | 11913 | 1612405 | 135 | 981 |
| M4 | 73954 | 11919231 | 161 | 982 |
| M5 | 33865 | 4518861 | 133 | 5875 |
| M6 | 33406 | 3200939 | 96 | 21510 |
| M7 | 56548 | 8190637 | 145 | 3950 |
| M8 | 15140 | 1805495 | 119 | 2401 |
| M9 | 42487 | 3055116 | 72 | 3300 |
| M10 | 29779 | 4226646 | 142 | 1200 |
| M11 | 35480 | 5892419 | 166 | 1566 |
| M12 | 30680 | 4830661 | 157 | 6729 |
| M13 | 19038 | 4491059 | 236 | 864 |
| M14 | 51683 | 8555982 | 166 | 10820 |
| M15 | - | - | - | 8407 |
| M16 | - | - | - | 4386 |
| M17 | 35789 | 5000270 | 140 | 4965 |
| M18 | 25894 | 4915988 | 190 | 2327 |
| M19 | 22086 | 3667640 | 166 | 236 |
| M20 | 66298 | 10545085 | 159 | 41178 |
| M21 | 76754 | 14431393 | 188 | - |
| M22 | 39398 | 6186789 | 157 | 813 |
| M23 | 69360 | 14053343 | 203 | 980 |
| M24 | - | - | - | 1690 |
| M25 | 236374 | 40447339 | 171 | - |
| M26 | - | - | - | 761 |

*M=Metastasis

**Supplemental Table 5. Multiplex Immunofluorescence IHC Antibodies**

| Antibody | Clone | Vendor | Species | Antigen Retrieval | Antibody Dilution Factor | Opal | Opal Dilution Factor | Opal Cat # |
|---|---|---|---|---|---|---|---|---|
| α-SMA | ASM$^{-}$1 | Leica | Mouse | EDTA, 20 min | RTU | 480 | 1:150 | FP1500<br>001KT |
| CD3 | LN10 | Leica | Mouse | EDTA, 20 min | RTU | 520 | 1:50 | FP1487<br>001KT |
| CD19 | BT51E | Leica | Mouse | EDTA, 20 min | RTU | 570 | 1:50 | FP1488<br>001KT |
| CD68 | 514H12 | Leica | Mouse | EDTA, 20 min | RTU | 620 | 1:50 | FP1495<br>001KT |
| IL-6 | 10C12 | Leica | Mouse | ETDA, 20 min | 1:500 | 690 | 1:50 | FP1497<br>001KT |
| CK19 | B170 | Leica | Mouse | ETDA, 20 min | RTU | 780 | 1:100 / 1:25 | FP1501<br>001KT |
| T-bet | EPR9301 | Abcam | Rabbit | EDTA, 20 min | 1:1000 | 480 | 1:150 | FP1500<br>001KT |
| RORγt | 6F3.1 | Biocare | Mouse | EDTA, 20 min | 1:100 | 520 | 1:50 | FP1487<br>001KT |
| CD4 | 4B12 | Leica | Mouse | EDTA, 20 min | RTU | 570 | 1:50 | FP1488<br>001KT |
| CD4 | SP35 | Cell Marque | Rabbit | EDTA, 20 min | 1:100 | 570 | 1:50 | FP1488<br>001Kt |
| CD8 | 4B11 | Leica | Mouse | EDTA, 20 min | RTU | 620 | 1:50 | FP1495<br>001KT |
| FoxP3 | 236A/E7 | Leica | Mouse | EDTA, 20 min | RTU | 690 | 1:50 | FP1497<br>001KT |
| DAPI | - | Akoya | - | - | - | - | - | - |

**Supplemental Table 6. Median of Minimum Distance Between Reference and Target Cells**

| Primary | | | Metastatic | | |
|---|---|---|---|---|---|
| Reference Cell | Target Cell | Minimum Distance (μm) | Reference Cell | Target Cell | Minimum Distance (μm) |
| $CK19^+$ | $CK19^+IL\text{-}6^+$ | 590.15 | $CK19^+$ | $CK19^+IL\text{-}6^+$ | 800.20 |
| | $\alpha\text{-}SMA^+$ | 265.42 | | $\alpha\text{-}SMA^+$ | 156.74 |
| | $\alpha\text{-}SMA^+IL\text{-}6^+$ | 2280.94 | | $\alpha\text{-}SMA^+IL\text{-}6^+$ | 2456.39 |
| | $CD3^+$ | 147.11 | | $CD3^+$ | 90.41 |
| | $CD3^+IL\text{-}6^+$ | 1997.61 | | $CD3^+IL\text{-}6^+$ | 1888.06 |
| | $CD19^+$ | 431.05 | | $CD19^+$ | 361.21 |
| | $CD19^+IL\text{-}6^+$ | 2197.58 | | $CD19^+IL\text{-}6^+$ | 383.74 |
| | $CD68^+$ | 74.54 | | $CD68^+$ | 47.25 |
| | $CD68^+IL\text{-}6^+$ | 1716.59 | | $CD68^+IL\text{-}6^+$ | 860.65 |
| $CK19^+IL\text{-}6^+$ | $CK19^+$ | 31.16 | $CK19^+IL\text{-}6^+$ | $CK19^+$ | 9.65 |
| | $\alpha\text{-}SMA^+$ | 201.31 | | $\alpha\text{-}SMA^+$ | 180.77 |
| | $\alpha\text{-}SMA^+IL\text{-}6^+$ | 1376.47 | | $\alpha\text{-}SMA^+IL\text{-}6^+$ | 1897.67 |
| | $CD3^+$ | 153.77 | | $CD3^+$ | 93.29 |
| | $CD3^+IL\text{-}6^+$ | 1035.86 | | $CD3^+IL\text{-}6^+$ | 1345.84 |
| | $CD19^+$ | 408.37 | | $CD19^+$ | 277.65 |
| | $CD19^+IL\text{-}6^+$ | 1659.47 | | $CD19^+IL\text{-}6^+$ | 604.41 |
| | $CD68^+$ | 85.24 | | $CD68^+$ | 53.66 |
| | $CD68^+IL\text{-}6^+$ | 831.24 | | $CD68^+IL\text{-}6^+$ | 594.47 |
| $\alpha\text{-}SMA^+$ | $CK19^+$ | 80.86 | $\alpha\text{-}SMA^+$ | $CK19^+$ | 24.49 |
| | $CK19^+IL\text{-}6^+$ | 580.02 | | $CK19^+IL\text{-}6^+$ | 778.60 |
| | $\alpha\text{-}SMA^+IL\text{-}6^+$ | 1492.05 | | $\alpha\text{-}SMA^+IL\text{-}6^+$ | 2561.07 |
| | $CD3^+$ | 143.35 | | $CD3^+$ | 82.68 |
| | $CD3^+IL\text{-}6^+$ | 1462.34 | | $CD3^+IL\text{-}6^+$ | 1758.43 |
| | $CD19^+$ | 392.82 | | $CD19^+$ | 430.39 |
| | $CD19^+IL\text{-}6^+$ | 1414.72 | | $CD19^+IL\text{-}6^+$ | 761.56 |
| | $CD68^+$ | 83.35 | | $CD68^+$ | 50.20 |
| | $CD68^+IL\text{-}6^+$ | 977.24 | | $CD68^+IL\text{-}6^+$ | 771.63 |
| $\alpha\text{-}SMA^+IL\text{-}6^+$ | $CK19^+$ | 107.29 | $\alpha\text{-}SMA^+IL\text{-}6^+$ | $CK19^+$ | 24.67 |
| | $CK19^+IL\text{-}6^+$ | 341.78 | | $CK19^+IL\text{-}6^+$ | 836.67 |
| | $\alpha\text{-}SMA^+$ | 111.84 | | $\alpha\text{-}SMA^+$ | 96.26 |
| | $CD3^+$ | 102.33 | | $CD3^+$ | 70.69 |
| | $CD3^+IL\text{-}6^+$ | 886.55 | | $CD3^+IL\text{-}6^+$ | 1732.74 |
| | $CD19^+$ | 649.53 | | $CD19^+$ | 387.19 |
| | $CD19^+IL\text{-}6^+$ | 1662.16 | | $CD19^+IL\text{-}6^+$ | 465.25 |
| | $CD68^+$ | 144.75 | | $CD68^+$ | 49.48 |
| | $CD68^+IL\text{-}6^+$ | 1050.49 | | $CD68^+IL\text{-}6^+$ | 569.52 |
| $CD3^+$ | $CK19^+$ | 74.10 | $CD3^+$ | $CK19^+$ | 34.57 |
| | $CK19^+IL\text{-}6^+$ | 674.41 | | $CK19^+IL\text{-}6^+$ | 863.28 |
| | $\alpha\text{-}SMA^+$ | 250.33 | | $\alpha\text{-}SMA^+$ | 149.14 |

| | | | | | |
|---|---|---|---|---|---|
| | α-SMA$^+$IL-6$^+$ | 2448.82 | | α-SMA$^+$IL-6$^+$ | 1178.00 |
| | CD3$^+$IL-6$^+$ | 1799.74 | | CD3$^+$IL-6$^+$ | 1233.66 |
| | CD19$^+$ | 226.05 | | CD19$^+$ | 211.67 |
| | CD19$^+$IL-6$^+$ | 1629.03 | | CD19$^+$IL-6$^+$ | 372.44 |
| | CD68$^+$ | 48.03 | | CD68$^+$ | 35.11 |
| | CD68$^+$IL-6$^+$ | 1188.42 | | CD68$^+$IL-6$^+$ | 1038.15 |
| CD3$^+$IL-6$^+$ | CK19$^+$ | 99.70 | CD3$^+$IL-6$^+$ | CK19$^+$ | 26.61 |
| | CK19$^+$IL-6$^+$ | 387.67 | | CK19$^+$IL-6$^+$ | 218.18 |
| | α-SMA$^+$ | 149.08 | | α-SMA$^+$ | 166.65 |
| | α-SMA$^+$IL-6$^+$ | 1454.78 | | α-SMA$^+$IL-6$^+$ | 1525.55 |
| | CD3$^+$ | 23.73 | | CD3$^+$ | 25.23 |
| | CD19$^+$ | 311.50 | | CD19$^+$ | 439.65 |
| | CD19$^+$IL-6$^+$ | 1008.21 | | CD19$^+$IL-6$^+$ | 864.19 |
| | CD68$^+$ | 89.91 | | CD68$^+$ | 46.04 |
| | CD68$^+$IL-6$^+$ | 492.29 | | CD68$^+$IL-6$^+$ | 482.62 |
| CD19$^+$ | CK19$^+$ | 111.60 | CD19$^+$ | CK19$^+$ | 34.02 |
| | CK19$^+$IL-6$^+$ | 599.28 | | CK19$^+$IL-6$^+$ | 1050.96 |
| | α-SMA$^+$ | 276.44 | | α-SMA$^+$ | 170.29 |
| | α-SMA$^+$IL-6$^+$ | 2202.85 | | α-SMA$^+$IL-6$^+$ | 1174.43 |
| | CD3$^+$ | 55.07 | | CD3$^+$ | 44.09 |
| | CD3$^+$IL-6$^+$ | 2027.00 | | CD3$^+$IL-6$^+$ | 1445.38 |
| | CD19$^+$IL-6$^+$ | 1661.84 | | CD19$^+$IL-6$^+$ | 422.57 |
| | CD68$^+$ | 68.39 | | CD68$^+$ | 41.15 |
| | CD68$^+$IL-6$^+$ | 1392.91 | | CD68$^+$IL-6$^+$ | 921.77 |
| CD19$^+$IL-6$^+$ | CK19$^+$ | 82.33 | CD19$^+$IL-6$^+$ | CK19$^+$ | 23.33 |
| | CK19$^+$IL-6$^+$ | 302.47 | | CK19$^+$IL-6$^+$ | 190.79 |
| | α-SMA$^+$ | 226.82 | | α-SMA$^+$ | 96.34 |
| | α-SMA$^+$IL-6$^+$ | 1927.55 | | α-SMA$^+$IL-6$^+$ | 1048.29 |
| | CD3$^+$ | 76.70 | | CD3$^+$ | 43.78 |
| | CD3$^+$IL-6$^+$ | 1030.24 | | CD3$^+$IL-6$^+$ | 997.46 |
| | CD19$^+$ | 44.16 | | CD19$^+$ | 94.58 |
| | CD68$^+$ | 103.68 | | CD68$^+$ | 41.95 |
| | CD68$^+$IL-6$^+$ | 598.40 | | CD68$^+$IL-6$^+$ | 521.77 |
| CD68$^+$ | CK19$^+$ | 69.98 | CD68$^+$ | CK19$^+$ | 31.11 |
| | CK19$^+$IL-6$^+$ | 677.57 | | CK19$^+$IL-6$^+$ | 1049.15 |
| | α-SMA$^+$ | 270.62 | | α-SMA$^+$ | 153.53 |
| | α-SMA$^+$IL-6$^+$ | 2238.78 | | α-SMA$^+$IL-6$^+$ | 1146.16 |
| | CD3$^+$ | 86.69 | | CD3$^+$ | 64.29 |
| | CD3$^+$IL-6$^+$ | 1585.88 | | CD3$^+$IL-6$^+$ | 1354.99 |
| | CD19$^+$ | 321.29 | | CD19$^+$ | 277.43 |
| | CD19$^+$IL-6$^+$ | 1717.99 | | CD19$^+$IL-6$^+$ | 580.39 |
| | CD68$^+$IL-6$^+$ | 1084.67 | | CD68$^+$IL-6$^+$ | 1110.30 |
| CD68$^+$IL-6$^+$ | CK19$^+$ | 51.95 | CD68$^+$IL-6$^+$ | CK19$^+$ | 14.99 |

| | | | | | |
|---|---|---|---|---|---|
| | CK19$^+$IL-6$^+$ | 186.73 | | CK19$^+$IL-6$^+$ | 281.10 |
| | α-SMA$^+$ | 150.11 | | α-SMA$^+$ | 104.29 |
| | α-SMA$^+$IL-6$^+$ | 1322.39 | | α-SMA$^+$IL-6$^+$ | 1729.94 |
| | CD3$^+$ | 72.86 | | CD3$^+$ | 116.60 |
| | CD3$^+$IL-6$^+$ | 700.81 | | CD3$^+$IL-6$^+$ | 617.58 |
| | CD19$^+$ | 309.77 | | CD19$^+$ | 449.44 |
| | CD19$^+$IL-6$^+$ | 1090.71 | | CD19$^+$IL-6$^+$ | 418.90 |
| | CD68$^+$ | 19.30 | | CD68$^+$ | 34.13 |

**Supplemental Table 7. Percent of Total Cells Phenotyped by mIHC in Each Defined Tumoral Structure**

| | % Phenotyped Cells in Tumor Structure | | | |
|---|---|---|---|---|
| **Sample** | **Tumoral** | **Internal** | **External** | **Stromal** |
| **P1** | - | 2.26 | 40.28 | 29.83 |
| **P3** | 17.03 | 14.59 | 45.44 | 38.97 |
| **P4** | 2.33 | 1.45 | 45.99 | 43.49 |
| **P5** | 4.29 | 2.47 | 44.95 | 39.30 |
| **P6** | 5.49 | 4.96 | 45.64 | 45.17 |
| **P8** | 9.32 | 4.96 | 46.07 | 44.93 |
| **P9** | 4.20 | 2.99 | 17.80 | 21.33 |
| **P10** | 9.54 | 8.69 | 46.35 | 44.81 |
| **P13** | 14.37 | 10.90 | 46.44 | 45.76 |
| **P14** | 17.93 | 13.10 | 46.39 | 44.28 |
| **P15** | 5.04 | 10.44 | 45.62 | 44.79 |
| **P16** | 2.99 | 12.31 | 43.55 | 41.84 |
| **P17** | 14.29 | 13.41 | 46.48 | 45.52 |
| **P18** | 7.02 | 5.17 | 46.27 | 39.20 |
| **P21** | 2.45 | 1.48 | 45.38 | 39.72 |
| **P22** | 7.16 | 5.64 | 46.25 | 42.50 |
| **P23** | 7.73 | 6.50 | 46.51 | 45.52 |
| **P24** | 6.04 | 8.29 | 46.44 | 45.80 |
| **P25** | 9.86 | 8.09 | 45.72 | 44.08 |
| **P27** | 8.51 | 9.06 | 39.69 | 33.53 |
| **M2** | - | 23.64 | 42.62 | 39.36 |
| **M3** | 9.33 | 8.23 | 39.46 | 19.52 |
| **M5** | 5.28 | 3.38 | 42.62 | 42.21 |
| **M8** | 8.96 | 4.67 | 37.16 | 13.81 |
| **M9** | 42.62 | 12.58 | 41.84 | 40.76 |
| **M12** | 6.84 | 5.94 | 42.62 | 41.92 |
| **M14** | 8.92 | 12.61 | 42.62 | 40.83 |
| **M19** | 18.19 | 16.94 | 42.41 | 41.22 |
| **M22** | 7.17 | 7.00 | 42.53 | 42.05 |
| **M23** | 4.00 | 4.15 | 42.52 | 39.09 |

**Supplemental Table 8. Percent of Each mIHC Phenotyped Cell Subtype in Each Defined Tumoral Structure**

| | Tumoral | | | | | | | |
|---|---|---|---|---|---|---|---|---|
| Sample | % CD19$^+$ | % CD19$^+$ IL-6$^+$ | % α-SMA$^+$ | % α-SMA$^+$ IL-6$^+$ | % CD68$^+$ | % CD68$^+$ IL-6$^+$ | % CD3$^+$ | % CD3$^+$ IL-6$^+$ |
| P1 | - | - | - | - | - | - | - | - |
| P3 | 0.74 | 0.00 | 78.22 | 0.50 | 6.44 | 0.00 | 14.11 | 0.00 |
| P4 | 11.69 | 0.00 | 53.37 | 0.00 | 4.09 | 0.00 | 30.84 | 0.00 |
| P5 | 2.29 | 0.00 | 75.24 | 0.00 | 8.55 | 0.00 | 13.91 | 0.00 |
| P6 | 1.09 | 0.00 | 59.93 | 0.00 | 1.34 | 0.00 | 37.64 | 0.00 |
| P8 | 1.52 | 0.00 | 56.75 | 0.00 | 0.14 | 0.00 | 41.59 | 0.00 |
| P9 | 92.31 | 0.00 | 0.00 | 0.00 | 0.00 | 0.00 | 7.69 | 0.00 |
| P10 | 2.04 | 0.00 | 64.11 | 0.00 | 2.42 | 0.00 | 31.43 | 0.00 |
| P13 | 0.30 | 0.00 | 54.51 | 0.00 | 16.08 | 0.00 | 29.12 | 0.00 |
| P14 | 2.06 | 0.06 | 0.53 | 0.00 | 0.06 | 0.00 | 97.24 | 0.06 |
| P15 | 0.00 | 0.00 | 5.97 | 0.00 | 2.99 | 1.49 | 89.55 | 0.00 |
| P16 | 11.84 | 0.00 | 1.32 | 0.00 | 1.32 | 0.00 | 85.53 | 0.00 |
| P17 | 0.46 | 0.00 | 72.37 | 0.00 | 1.29 | 0.00 | 25.88 | 0.00 |
| P18 | 2.42 | 0.00 | 3.49 | 0.00 | 3.32 | 0.00 | 90.77 | 0.00 |
| P21 | 0.74 | 0.00 | 5.45 | 0.00 | 4.21 | 0.00 | 89.60 | 0.00 |
| P22 | 0.37 | 0.00 | 4.97 | 0.00 | 0.82 | 0.00 | 93.84 | 0.00 |
| P23 | 1.64 | 0.00 | 5.39 | 0.00 | 0.95 | 0.00 | 91.89 | 0.14 |
| P24 | 12.67 | 0.10 | 73.42 | 0.03 | 0.03 | 0.00 | 13.74 | 0.01 |
| P25 | 1.64 | 0.00 | 2.62 | 0.00 | 5.90 | 0.00 | 89.84 | 0.00 |
| P27 | 0.00 | 0.00 | 8.70 | 13.04 | 0.00 | 0.00 | 39.13 | 39.13 |
| M2 | - | - | - | - | - | - | - | - |
| M3 | 2.27 | 0.00 | 2.27 | 0.00 | 0.00 | 0.00 | 86.36 | 9.09 |
| M5 | 6.44 | 0.00 | 45.45 | 0.00 | 0.00 | 0.00 | 47.94 | 0.17 |
| M8 | 5.17 | 0.00 | 1.72 | 0.00 | 6.90 | 3.45 | 60.34 | 22.41 |
| M9 | 0.00 | 0.00 | 100.00 | 0.00 | 0.00 | 0.00 | 0.00 | 0.00 |
| M12 | 1.56 | 0.00 | 15.11 | 0.00 | 8.03 | 0.00 | 75.31 | 0.00 |
| M14 | 1.56 | 0.00 | 56.33 | 0.00 | 3.88 | 0.00 | 38.23 | 0.00 |
| M19 | 0.15 | 0.00 | 19.85 | 0.00 | 1.31 | 0.00 | 78.69 | 0.00 |
| M22 | 61.96 | 0.00 | 6.18 | 0.00 | 0.15 | 0.00 | 31.72 | 0.00 |
| M23 | 1.04 | 0.00 | 21.43 | 0.00 | 8.18 | 0.00 | 69.35 | 0.00 |

| | Internal Margin | | | | | | | |
|---|---|---|---|---|---|---|---|---|
| Sample | % CD19$^+$ | % CD19$^+$ IL-6$^+$ | % α-SMA$^+$ | % α-SMA$^+$ IL-6$^+$ | % CD68$^+$ | % CD68$^+$ IL-6$^+$ | % CD3$^+$ | % CD3$^+$ IL-6$^+$ |
| P1 | 29.63 | 0.00 | 22.22 | 0.00 | 7.41 | 3.70 | 33.33 | 3.70 |
| P3 | 0.62 | 0.00 | 74.10 | 0.62 | 10.37 | 0.08 | 14.20 | 0.00 |
| P4 | 15.13 | 0.00 | 41.06 | 0.00 | 3.28 | 0.00 | 40.52 | 0.00 |
| P5 | 4.91 | 0.00 | 66.35 | 0.00 | 12.25 | 0.00 | 16.49 | 0.00 |
| P6 | 1.80 | 0.00 | 54.22 | 0.00 | 4.79 | 0.00 | 39.20 | 0.00 |
| P8 | 3.18 | 0.00 | 53.99 | 0.00 | 0.32 | 0.00 | 42.51 | 0.00 |

| | | | | | | | | |
|---|---|---|---|---|---|---|---|---|
| **P9** | 76.60 | 0.00 | 0.00 | 0.00 | 0.00 | 0.00 | 23.40 | 0.00 |
| **P10** | 6.79 | 0.00 | 56.74 | 0.00 | 5.67 | 0.00 | 30.80 | 0.00 |
| **P13** | 1.91 | 0.00 | 50.09 | 0.00 | 15.08 | 0.00 | 32.76 | 0.17 |
| **P14** | 2.29 | 0.00 | 0.90 | 0.00 | 0.14 | 0.00 | 96.60 | 0.07 |
| **P15** | 1.31 | 0.44 | 13.35 | 0.00 | 11.60 | 0.66 | 70.68 | 1.97 |
| **P16** | 1.40 | 0.00 | 22.46 | 0.00 | 26.14 | 0.00 | 50.00 | 0.00 |
| **P17** | 0.92 | 0.05 | 72.22 | 0.00 | 1.24 | 0.00 | 25.56 | 0.00 |
| **P18** | 2.02 | 0.00 | 2.14 | 0.12 | 4.52 | 0.12 | 90.84 | 0.24 |
| **P21** | 4.93 | 0.00 | 1.48 | 0.00 | 0.00 | 0.00 | 92.61 | 0.99 |
| **P22** | 1.86 | 0.00 | 3.58 | 0.00 | 1.72 | 0.00 | 92.85 | 0.00 |
| **P23** | 3.01 | 0.00 | 3.16 | 0.00 | 0.45 | 0.00 | 93.38 | 0.00 |
| **P24** | 13.74 | 0.13 | 75.85 | 0.06 | 0.04 | 0.00 | 10.18 | 0.00 |
| **P25** | 1.63 | 0.41 | 5.84 | 0.00 | 5.98 | 0.27 | 85.33 | 0.54 |
| **P27** | 2.04 | 0.00 | 10.20 | 4.08 | 0.00 | 0.00 | 56.12 | 27.55 |
| **M2** | 12.68 | 1.41 | 77.46 | 2.82 | 0.00 | 0.00 | 5.63 | 0.00 |
| **M3** | 13.64 | 1.52 | 6.06 | 0.00 | 0.00 | 0.00 | 74.24 | 4.55 |
| **M5** | 12.86 | 0.00 | 44.76 | 0.00 | 0.00 | 0.00 | 41.90 | 0.48 |
| **M8** | 7.27 | 0.00 | 1.82 | 0.00 | 9.09 | 14.55 | 47.27 | 20.00 |
| **M9** | 1.11 | 0.00 | 72.22 | 0.00 | 1.11 | 0.00 | 25.56 | 0.00 |
| **M12** | 3.45 | 0.00 | 12.56 | 0.00 | 5.42 | 0.00 | 78.57 | 0.00 |
| **M14** | 2.57 | 0.00 | 63.88 | 0.00 | 1.12 | 0.00 | 32.26 | 0.16 |
| **M19** | 0.33 | 0.00 | 22.17 | 0.00 | 0.99 | 0.00 | 76.52 | 0.00 |
| **M22** | 58.42 | 0.00 | 6.03 | 0.00 | 0.21 | 0.00 | 35.34 | 0.00 |
| **M23** | 2.89 | 0.00 | 29.11 | 0.00 | 3.56 | 0.00 | 64.44 | 0.00 |

| | **External Margin** | | | | | | | |
|---|---|---|---|---|---|---|---|---|
| **Sample** | **% $CD19^+$** | **% $CD19^+$ $IL\text{-}6^+$** | **% $\alpha\text{-}SMA^+$** | **% $\alpha\text{-}SMA^+$ $IL\text{-}6^+$** | **% $CD68^+$** | **% $CD68^+$ $IL\text{-}6^+$** | **% $CD3^+$** | **% $CD3^+$ $IL\text{-}6^+$** |
| **P1** | 13.10 | 0.00 | 52.38 | 2.38 | 14.29 | 0.00 | 17.86 | 0.00 |
| **P3** | 0.63 | 0.00 | 84.40 | 0.63 | 6.58 | 0.18 | 7.57 | 0.00 |
| **P4** | 8.13 | 0.00 | 49.44 | 0.00 | 5.48 | 0.00 | 36.94 | 0.00 |
| **P5** | 4.92 | 0.00 | 89.07 | 0.00 | 2.11 | 0.00 | 3.90 | 0.00 |
| **P6** | 1.45 | 0.00 | 65.88 | 0.00 | 3.21 | 0.00 | 29.47 | 0.00 |
| **P8** | 1.71 | 0.00 | 66.19 | 0.00 | 0.53 | 0.00 | 31.56 | 0.00 |
| **P9** | 94.08 | 0.00 | 3.55 | 0.00 | 0.00 | 0.00 | 2.37 | 0.00 |
| **P10** | 8.70 | 0.00 | 55.02 | 0.00 | 6.02 | 0.00 | 30.27 | 0.00 |
| **P13** | 2.40 | 0.00 | 54.60 | 0.00 | 17.26 | 0.00 | 25.74 | 0.00 |
| **P14** | 4.02 | 0.00 | 0.94 | 0.00 | 0.26 | 0.00 | 94.69 | 0.09 |
| **P15** | 0.88 | 0.00 | 12.04 | 0.10 | 12.72 | 0.20 | 73.78 | 0.29 |
| **P16** | 4.89 | 0.00 | 13.04 | 0.00 | 11.96 | 0.82 | 69.29 | 0.00 |
| **P17** | 0.93 | 0.00 | 74.13 | 0.00 | 1.63 | 0.00 | 23.31 | 0.00 |
| **P18** | 3.66 | 0.10 | 2.54 | 0.00 | 7.01 | 0.10 | 86.38 | 0.20 |
| **P21** | 3.74 | 0.00 | 4.98 | 0.00 | 4.05 | 0.31 | 86.92 | 0.00 |
| **P22** | 3.63 | 0.00 | 5.87 | 0.00 | 1.12 | 0.00 | 89.39 | 0.00 |

| | | | | | | | | |
|---|---|---|---|---|---|---|---|---|
| **P23** | 2.27 | 0.00 | 3.07 | 0.00 | 0.53 | 0.00 | 94.13 | 0.00 |
| **P24** | 13.29 | 0.16 | 75.89 | 0.02 | 0.35 | 0.00 | 10.29 | 0.02 |
| **P25** | 2.21 | 0.25 | 15.81 | 0.12 | 5.88 | 0.00 | 75.74 | 0.00 |
| **P27** | 3.03 | 3.03 | 39.39 | 18.18 | 0.00 | 1.01 | 28.28 | 7.07 |
| **M2** | 28.43 | 0.00 | 53.92 | 0.98 | 0.00 | 0.00 | 15.69 | 0.98 |
| **M3** | 24.00 | 0.00 | 16.00 | 0.00 | 0.00 | 0.00 | 60.00 | 0.00 |
| **M5** | 17.86 | 0.00 | 58.33 | 0.00 | 0.00 | 0.00 | 23.81 | 0.00 |
| **M8** | 8.82 | 5.88 | 11.76 | 0.00 | 17.65 | 11.76 | 35.29 | 8.82 |
| **M9** | 1.49 | 0.00 | 69.52 | 0.00 | 0.74 | 0.00 | 28.25 | 0.00 |
| **M12** | 2.33 | 0.00 | 18.60 | 0.00 | 6.98 | 0.00 | 72.09 | 0.00 |
| **M14** | 0.27 | 0.00 | 52.68 | 0.00 | 0.54 | 0.00 | 46.51 | 0.00 |
| **M19** | 0.67 | 0.00 | 30.95 | 0.00 | 1.33 | 0.00 | 66.22 | 0.83 |
| **M22** | 63.08 | 0.00 | 9.01 | 0.00 | 0.00 | 0.00 | 27.91 | 0.00 |
| **M23** | 2.73 | 0.00 | 37.59 | 0.00 | 5.24 | 0.00 | 54.44 | 0.00 |

| | **Stroma** | | | | | | | |
|---|---|---|---|---|---|---|---|---|
| **Sample** | **% CD19$^+$** | **% CD19$^+$ IL-6$^+$** | **% α-SMA$^+$** | **% α-SMA$^+$ IL-6$^+$** | **% CD68$^+$** | **% CD68$^+$ IL-6$^+$** | **% CD3$^+$** | **% CD3$^+$ IL-6$^+$** |
| **P1** | 8.99 | 0.21 | 27.84 | 1.71 | 4.50 | 1.28 | 52.46 | 3.00 |
| **P3** | 0.34 | 0.03 | 95.14 | 0.38 | 1.67 | 0.06 | 2.40 | 0.00 |
| **P4** | 10.53 | 0.00 | 54.07 | 0.00 | 15.18 | 0.00 | 20.22 | 0.00 |
| **P5** | 16.88 | 0.00 | 71.64 | 0.00 | 7.52 | 0.00 | 3.96 | 0.00 |
| **P6** | 2.63 | 0.00 | 61.01 | 0.00 | 4.42 | 0.00 | 31.93 | 0.00 |
| **P8** | 4.34 | 0.00 | 76.66 | 0.00 | 1.33 | 0.00 | 17.67 | 0.00 |
| **P9** | 82.77 | 0.00 | 13.92 | 0.00 | 0.27 | 0.00 | 3.04 | 0.00 |
| **P10** | 11.84 | 0.00 | 62.13 | 0.00 | 3.50 | 0.00 | 22.54 | 0.00 |
| **P13** | 3.47 | 0.00 | 50.61 | 0.03 | 14.35 | 0.00 | 31.53 | 0.00 |
| **P14** | 3.90 | 0.00 | 0.53 | 0.03 | 0.30 | 0.00 | 95.20 | 0.03 |
| **P15** | 1.01 | 0.00 | 5.93 | 0.00 | 2.34 | 0.02 | 90.54 | 0.15 |
| **P16** | 2.22 | 0.03 | 9.54 | 0.03 | 23.17 | 0.00 | 64.88 | 0.13 |
| **P17** | 0.87 | 0.01 | 72.27 | 0.00 | 4.40 | 0.00 | 22.44 | 0.01 |
| **P18** | 6.32 | 0.04 | 2.77 | 0.09 | 17.84 | 0.26 | 72.42 | 0.26 |
| **P21** | 0.56 | 0.09 | 7.51 | 0.00 | 3.94 | 0.00 | 87.61 | 0.28 |
| **P22** | 3.20 | 0.00 | 9.36 | 0.00 | 0.86 | 0.00 | 86.58 | 0.00 |
| **P23** | 0.72 | 0.00 | 2.78 | 0.02 | 2.57 | 0.00 | 93.78 | 0.13 |
| **P24** | 14.11 | 0.11 | 71.57 | 0.03 | 0.75 | 0.00 | 13.41 | 0.02 |
| **P25** | 1.11 | 0.03 | 19.94 | 0.03 | 24.85 | 0.15 | 53.75 | 0.13 |
| **P27** | 13.28 | 4.80 | 32.77 | 16.67 | 0.00 | 0.00 | 20.62 | 11.86 |
| **M2** | 21.66 | 0.00 | 57.32 | 1.91 | 5.10 | 0.00 | 14.01 | 0.00 |
| **M3** | 16.67 | 0.00 | 10.00 | 0.00 | 0.00 | 0.00 | 68.33 | 5.00 |
| **M5** | 2.94 | 0.00 | 82.35 | 0.00 | 0.00 | 0.00 | 14.71 | 0.00 |
| **M8** | 24.14 | 1.72 | 5.17 | 1.72 | 25.86 | 10.34 | 25.86 | 5.17 |
| **M9** | 2.57 | 0.04 | 35.87 | 0.04 | 0.17 | 0.04 | 61.02 | 0.25 |
| **M12** | 2.50 | 0.00 | 42.08 | 0.00 | 9.17 | 0.00 | 46.25 | 0.00 |

| | | | | | | | | |
|---|---|---|---|---|---|---|---|---|
| **M14** | 0.61 | 0.00 | 27.38 | 0.00 | 0.90 | 0.00 | 71.12 | 0.00 |
| **M19** | 0.32 | 0.00 | 52.26 | 0.05 | 2.39 | 0.00 | 44.98 | 0.00 |
| **M22** | 57.44 | 0.00 | 11.04 | 0.00 | 0.08 | 0.00 | 31.44 | 0.00 |
| **M23** | 1.71 | 0.00 | 41.06 | 0.00 | 11.54 | 0.00 | 45.69 | 0.00 |

**Supplemental Table 9. CyTOF Antibody Panel**

| CyTOF Antibodies | | |
|---|---|---|
| **Marker** | **Clone** | **Metal** |
| CD45 | HI30 | 89Y |
| B7-H3 | MIH42 | 141Pr |
| CD19 | HIB19 | 142Nd |
| HLA-DR | L243 | 143Nd |
| CD69 | FN50 | 144Nd |
| CD4 | RPA-T4 | 145Nd |
| CD8 | RPA-T8 | 146Nd |
| ICOS | C398.4A | 148Nd |
| CD127 | A019D5 | 149Sm |
| CD103 | Ber-ACT8 | 151Eu |
| CD95 | DX9 | 152Sm |
| TIGIT | MBSA43 | 153Eu |
| TIM-3 | F38-2E2 | 154Sm |
| CD27 | L128 | 155Gd |
| PD-L1 | 29E.2A3 | 156Gd |
| CD33 | WM53 | 158Gd |
| CCR7 | G043H7 | 159Tb |
| CXCR5 | 51505 | 164Dy |
| CD45RO | UCHL1 | 165Ho |
| NKG2D | ON72 | 166Er |
| CD11b | ICRF44 | 167Er |
| CD25 | 2A3 | 169Tm |
| CD3 | SP34-2 | 170Er |
| CD38 | HIT2 | 172Yb |
| CXCR4 | 12G5 | 173Yb |
| PD-1 | EH12.2H7 | 174Yb |
| CD14 | M5E2 | 175Lu |
| CD56 | CMSSB | 176Yb |
| CD16 | 3G8 | 209Bi |
| TCF-1 | 7F11.A10 | 147Sm |
| LAG-3 | 11C3C65 | 150Nd |
| T-Bet | 4B10 | 160Gd |
| CTLA-4 | 14D3 | 161Dy |
| FoxP3 | PCH101 | 162Dy |
| EOMES | WD1928 | 163Dy |
| Ki-67 | B56 | 168Er |
| Granzyme B | CB11 | 171Yb |

**Supplemental Table 10. Immune Cell Populations**

| Immune Cells | Primary (n=12) | | | Metastatic (n=23) | | |
|---|---|---|---|---|---|---|
| | **Mean** | **SD** | **Median** | **Mean** | **SD** | **Median** |
| **Live Single Cells** | | | | | | |
| $CD45^-$ | 63.78% | 35.25% | 74.25% | 66.76% | 36.79% | 91.50% |
| $CD45^+$ | 36.23% | 35.24% | 25.75% | 33.25% | 36.79% | 8.50% |
| **Lineage - % Live Cells** | | | | | | |
| $CD3^+$ | 11.63% | 18.69% | 2.14% | 15.73% | 22.37% | 1.66% |
| $CD4^+$ | 7.45% | 12.86% | 0.52% | 5.80% | 8.69% | 0.64% |
| $CD8^+$ | 3.55% | 5.29% | 1.00% | 8.35% | 12.34% | 0.96% |
| $CD19^+$ | 1.83% | 3.57% | 0.38% | 1.88% | 3.69% | 0.76% |
| $CD11b^+$ | 20.07% | 31.52% | 6.61% | 7.44% | 10.06% | 1.92% |
| $CD56^+$ $CD3^-$ | 0.35% | 0.40% | 0.19% | 0.18% | 0.45% | 0.01% |
| **Ki-67** | | | | | | |
| $CD4^+$ T Cells | 6.09% | 6.64% | 5.33% | 8.86% | 10.39% | 5.69% |
| $CD8^+$ T Cells | 9.23% | 12.02% | 4.57% | 11.54% | 8.38% | 8.78% |
| $CD19^+$ Cells | 12.02% | 18.68% | 3.23% | 12.12% | 8.01% | 12.30% |

**Supplemental Table 11. T Cell Counts**

| Samples Used for T Cell Subset Analysis | | | |
|---|---|---|---|
| **Sample** | **Total T Cells** | **Total CD4$^+$** | **Total CD8$^+$** |
| P7 | 7761 | 5036 | 2323 |
| P8 | 1904 | 1419 | 448 |
| P11 | 710 | 447 | 238 |
| P20 | 25 | 11 | 13 |
| P22 | 27 | 10 | 11 |
| P23 | 33 | 6 | 13 |
| P26 | 159 | 97 | 52 |
| P27 | 70 | 38 | 27 |
| M8 | 772 | 387 | 318 |
| M10 | 42 | 25 | 15 |
| M11 | 26 | 8 | 15 |
| M12 | 2975 | 816 | 1913 |
| M13 | 138 | 59 | 67 |
| M14 | 7514 | 2308 | 4153 |
| M15 | 3594 | 1663 | 1557 |
| M16 | 1974 | 351 | 1455 |
| M17 | 60 | 32 | 22 |
| M18 | 1498 | 720 | 673 |
| M23 | 201 | 100 | 88 |
| M24 | 132 | 49 | 79 |
| M26 | 87 | 13 | 54 |
| **Samples Excluded Due to T Cell Number** | | | |
| **Sample** | **Total T Cells** | **Total CD4$^+$** | **Total CD8$^+$** |
| P9 | 1 | 1 | 0 |
| P10 | 24 | 16 | 7 |
| P21 | 18 | 4 | 11 |
| P25 | 2 | 0 | 0 |
| M1 | 4 | 0 | 1 |
| M2 | 0 | 0 | 0 |
| M3 | 2 | 1 | 0 |
| M4 | 3 | 0 | 0 |
| M5 | 3 | 0 | 0 |
| M6 | 3 | 1 | 1 |
| M7 | 1 | 1 | 0 |
| M9 | 9 | 5 | 3 |
| M20 | 1 | 1 | 0 |
| M22 | 2 | 1 | 0 |

**Supplemental Table 12. CD11b$^+$ Cell Counts**

| Samples Used for CD11b$^+$ Subset Analysis | |
|---|---|
| **Sample** | **Total CD11b$^+$** |
| P7 | 2877 |
| P8 | 328 |
| P11 | 474 |
| P22 | 31 |
| P23 | 144 |
| P26 | 26503 |
| P27 | 16094 |
| M5 | 73 |
| M6 | 33 |
| M7 | 76 |
| M8 | 212 |
| M9 | 94 |
| M10 | 26 |
| M12 | 1719 |
| M13 | 233 |
| M14 | 628 |
| M15 | 902 |
| M16 | 669 |
| M18 | 383 |
| M23 | 336 |
| M26 | 104 |
| **Samples Excluded Due to CD11b$^+$ Cell Number** | |
| **Sample** | **Total CD11b$^+$** |
| P9 | 4 |
| P10 | 19 |
| P20 | 18 |
| P21 | 3 |
| P25 | 0 |
| M1 | 4 |
| M2 | 6 |
| M3 | 7 |
| M4 | 15 |
| M11 | 3 |
| M17 | 8 |
| M20 | 10 |
| M22 | 6 |
| M24 | 16 |

**Supplemental Table 13. CD19$^+$ Cell Counts**

| Samples Used for CD19$^+$ Subset Analysis | |
|---|---|
| **Sample** | **Total CD19$^+$** |
| P7 | 213 |
| P11 | 111 |
| P22 | 62 |
| P26 | 25 |
| M2 | 76 |
| M4 | 28 |
| M5 | 45 |
| M7 | 69 |
| M8 | 219 |
| M9 | 537 |
| M12 | 69 |
| M14 | 82 |
| M15 | 80 |
| M16 | 30 |
| **Samples Excluded Due to CD19$^+$ Cell Number** | |
| **Sample** | **Total CD19$^+$** |
| P8 | 21 |
| P9 | 4 |
| P10 | 11 |
| P20 | 10 |
| P21 | 5 |
| P23 | 3 |
| P25 | 1 |
| P27 | 3 |
| M1 | 3 |
| M3 | 10 |
| M6 | 2 |
| M10 | 7 |
| M11 | 12 |
| M13 | 9 |
| M17 | 7 |
| M18 | 7 |
| M20 | 12 |
| M22 | 3 |
| M23 | 3 |
| M24 | 1 |
| M26 | 2 |

**Supplemental Table 14. CD4$^{+}$ T Cell Checkpoint Combination Frequency**

| Immune Cells | Primary (n=8) | | | Metastatic (n=13) | | |
|---|---|---|---|---|---|---|
| | Mean | SD | Median | Mean | SD | Median |
| **Total CD4$^{+}$** | | | | | | |
| CTLA-4 | 85.08% | 15.41% | 87.80% | 87.25% | 9.61% | 88.30% |
| LAG-3 | 78.90% | 24.30% | 88.95% | 76.08% | 15.24% | 75.00% |
| PD-1 | 75.79% | 12.53% | 73.75% | 76.15% | 14.55% | 81.80% |
| TIGIT | 41.26% | 12.61% | 45.95% | 38.38% | 14.00% | 37.30% |
| TIM-3 | 31.98% | 15.38% | 30.45% | 28.54% | 16.88% | 22.20% |
| **0 Checkpoints** | | | | | | |
| CTLA-4$^{-}$LAG-3$^{-}$PD-1$^{-}$TIGIT$^{-}$TIM-3$^{-}$ | 0.45% | 0.78% | 0.00% | 0.18% | 0.34% | 0.00% |
| **1 Checkpoint** | | | | | | |
| CTLA-4$^{+}$LAG-3$^{-}$PD-1$^{-}$TIGIT$^{-}$TIM-3$^{-}$ | 2.94% | 5.74% | 0.39% | 3.73% | 3.74% | 2.64% |
| CTLA-4$^{-}$LAG-3$^{+}$PD-1$^{-}$TIGIT$^{-}$TIM-3$^{-}$ | 0.50% | 0.77% | 0.00% | 1.92% | 3.44% | 1.02% |
| CTLA-4$^{-}$LAG-3$^{-}$PD-1$^{+}$TIGIT$^{-}$TIM-3$^{-}$ | 0.49% | 0.85% | 0.11% | 1.03% | 1.35% | 0.00% |
| CTLA-4$^{-}$LAG-3$^{-}$PD-1$^{-}$TIGIT$^{+}$TIM-3$^{-}$ | 0.02% | 0.03% | 0.00% | 0.08% | 0.28% | 0.00% |
| CTLA-4$^{-}$LAG-3$^{-}$PD-1$^{-}$TIGIT$^{-}$TIM-3$^{+}$ | 0.02% | 0.05% | 0.00% | 0.06% | 0.11% | 0.00% |
| **2 Checkpoints** | | | | | | |
| CTLA-4$^{+}$LAG-3$^{+}$PD-1$^{-}$TIGIT$^{-}$TIM-3$^{-}$ | 7.05% | 6.61% | 6.58% | 8.35% | 7.29% | 6.78% |
| CTLA-4$^{+}$LAG-3$^{-}$PD-1$^{+}$TIGIT$^{-}$TIM-3$^{-}$ | 6.35% | 7.47% | 3.29% | 7.98% | 6.41% | 6.94% |
| CTLA-4$^{+}$LAG-3$^{-}$PD-1$^{-}$TIGIT$^{+}$TIM-3$^{-}$ | 0.27% | 0.37% | 0.11% | 0.21% | 0.32% | 0.00% |
| CTLA-4$^{+}$LAG-3$^{-}$PD-1$^{-}$TIGIT$^{-}$TIM-3$^{+}$ | 0.47% | 0.81% | 0.00% | 0.78% | 1.16% | 0.30% |
| CTLA-4$^{-}$LAG-3$^{+}$PD-1$^{+}$TIGIT$^{-}$TIM-3$^{-}$ | 2.09% | 3.46% | 0.34% | 3.10% | 4.25% | 1.69% |
| CTLA-4$^{-}$LAG-3$^{+}$PD-1$^{-}$TIGIT$^{+}$TIM-3$^{-}$ | 1.43% | 3.48% | 0.07% | 0.07% | 0.13% | 0.00% |
| CTLA-4$^{-}$LAG-3$^{+}$PD-1$^{-}$TIGIT$^{-}$TIM-3$^{+}$ | 0.20% | 0.33% | 0.00% | 0.37% | 0.85% | 0.00% |
| CTLA-4$^{-}$LAG-3$^{-}$PD-1$^{+}$TIGIT$^{+}$TIM-3$^{-}$ | 0.13% | 0.20% | 0.00% | 1.37% | 3.52% | 0.00% |
| CTLA-4$^{-}$LAG-3$^{-}$PD-1$^{+}$TIGIT$^{-}$TIM-3$^{+}$ | 0.07% | 0.15% | 0.00% | 0.21% | 0.47% | 0.00% |
| CTLA-4$^{-}$LAG-3$^{-}$PD-1$^{-}$TIGIT$^{+}$TIM-3$^{+}$ | 0.02% | 0.05% | 0.00% | 0.01% | 0.04% | 0.00% |
| **3 Checkpoints** | | | | | | |
| CTLA-4$^{+}$LAG-3$^{+}$PD-1$^{+}$TIGIT$^{-}$TIM-3$^{-}$ | 18.71% | 11.10% | 18.05% | 20.22% | 11.42% | 23.10% |
| CTLA-4$^{+}$LAG-3$^{+}$PD-1$^{-}$TIGIT$^{+}$TIM-3$^{-}$ | 3.58% | 3.63% | 2.86% | 2.44% | 3.49% | 1.59% |
| CTLA-4$^{+}$LAG-3$^{+}$PD-1$^{-}$TIGIT$^{-}$TIM-3$^{+}$ | 6.16% | 5.73% | 5.65% | 4.21% | 9.23% | 2.04% |
| CTLA-4$^{+}$LAG-3$^{-}$PD-1$^{+}$TIGIT$^{+}$TIM-3$^{-}$ | 6.70% | 11.12% | 2.40% | 3.52% | 4.40% | 2.29% |
| CTLA-4$^{+}$LAG-3$^{-}$PD-1$^{+}$TIGIT$^{-}$TIM-3$^{+}$ | 1.18% | 2.02% | 0.30% | 2.24% | 3.49% | 0.91% |
| CTLA-4$^{+}$LAG-3$^{-}$PD-1$^{-}$TIGIT$^{+}$TIM-3$^{+}$ | 0.11% | 0.23% | 0.00% | 0.05% | 0.10% | 0.00% |
| CTLA-4$^{-}$LAG-3$^{+}$PD-1$^{+}$TIGIT$^{+}$TIM-3$^{-}$ | 4.09% | 6.08% | 1.57% | 2.29% | 3.38% | 1.69% |
| CTLA-4$^{-}$LAG-3$^{+}$PD-1$^{+}$TIGIT$^{-}$TIM-3$^{+}$ | 3.91% | 7.20% | 0.50% | 1.30% | 2.34% | 0.37% |
| CTLA-4$^{-}$LAG-3$^{+}$PD-1$^{-}$TIGIT$^{+}$TIM-3$^{+}$ | 0.03% | 0.07% | 0.00% | 0.01% | 0.05% | 0.00% |
| CTLA-4$^{-}$LAG-3$^{-}$PD-1$^{+}$TIGIT$^{+}$TIM-3$^{+}$ | 0.05% | 0.09% | 0.00% | 0.24% | 0.86% | 0.00% |
| **4 Checkpoints** | | | | | | |
| CTLA-4$^{+}$LAG-3$^{+}$PD-1$^{+}$TIGIT$^{+}$TIM-3$^{-}$ | 13.26% | 11.58% | 12.75% | 14.97% | 8.33% | 14.90% |
| CTLA-4$^{+}$LAG-3$^{+}$PD-1$^{+}$TIGIT$^{-}$TIM-3$^{+}$ | 8.17% | 8.78% | 7.82% | 5.94% | 3.23% | 6.12% |
| CTLA-4$^{+}$LAG-3$^{+}$PD-1$^{-}$TIGIT$^{+}$TIM-3$^{+}$ | 0.97% | 1.69% | 0.00% | 1.37% | 2.23% | 0.52% |

| | | | | | | |
|---|---|---|---|---|---|---|
| CTLA-4$^+$LAG-3$^-$PD-1$^+$TIGIT$^+$TIM-3$^+$ | 1.82% | 2.03% | 1.25% | 2.21% | 3.33% | 0.69% |
| CTLA-4$^-$LAG-3$^+$PD-1$^+$TIGIT$^+$TIM-3$^+$ | 1.44% | 3.13% | 0.07% | 0.48% | 0.68% | 0.25% |
| **5 Checkpoints** | | | | | | |
| CTLA-4$^+$LAG-3$^+$PD-1$^+$TIGIT$^+$TIM-3$^+$ | 7.36% | 5.60% | 8.49% | 9.04% | 5.41% | 7.75% |

**Supplemental Table 15. CD8$^{+}$ T Cell Checkpoint Combination Frequency**

| Immune Cells | Primary (n=8) | | | Metastatic (n=13) | | |
|---|---|---|---|---|---|---|
| | Mean | SD | Median | Mean | SD | Median |
| **Total CD8$^{+}$** | | | | | | |
| CTLA-4 | 42.06% | 26.61% | 44.75% | 60.07% | 17.20% | 54.60% |
| LAG-3 | 57.03% | 35.39% | 56.25% | 44.72% | 28.30% | 29.90% |
| PD-1 | 73.31% | 18.39% | 75.35% | 71.95% | 17.68% | 76.10% |
| TIGIT | 50.61% | 20.93% | 49.85% | 62.84% | 15.69% | 59.10% |
| TIM-3 | 25.45% | 13.29% | 23.50% | 20.01% | 14.50% | 15.90% |
| **0 Checkpoints** | | | | | | |
| CTLA-4$^{-}$LAG-3$^{-}$PD-1$^{-}$TIGIT$^{-}$TIM-3$^{-}$ | 5.57% | 10.61% | 1.17% | 1.69% | 1.95% | 1.49% |
| **1 Checkpoint** | | | | | | |
| CTLA-4$^{+}$LAG-3$^{-}$PD-1$^{-}$TIGIT$^{-}$TIM-3$^{-}$ | 1.41% | 1.75% | 0.84% | 2.43% | 2.29% | 2.06% |
| CTLA-4$^{-}$LAG-3$^{+}$PD-1$^{-}$TIGIT$^{-}$TIM-3$^{-}$ | 4.17% | 6.22% | 2.17% | 1.19% | 1.11% | 1.49% |
| CTLA-4$^{-}$LAG-3$^{-}$PD-1$^{+}$TIGIT$^{-}$TIM-3$^{-}$ | 4.24% | 4.71% | 3.51% | 5.23% | 4.23% | 6.21% |
| CTLA-4$^{-}$LAG-3$^{-}$PD-1$^{-}$TIGIT$^{+}$TIM-3$^{-}$ | 2.15% | 2.84% | 0.63% | 3.51% | 4.53% | 1.04% |
| CTLA-4$^{-}$LAG-3$^{-}$PD-1$^{-}$TIGIT$^{-}$TIM-3$^{+}$ | 1.53% | 2.63% | 0.30% | 0.31% | 0.58% | 0.00% |
| **2 Checkpoints** | | | | | | |
| CTLA-4$^{+}$LAG-3$^{+}$PD-1$^{-}$TIGIT$^{-}$TIM-3$^{-}$ | 2.34% | 4.07% | 0.45% | 3.58% | 7.75% | 1.09% |
| CTLA-4$^{+}$LAG-3$^{-}$PD-1$^{+}$TIGIT$^{-}$TIM-3$^{-}$ | 4.24% | 5.40% | 1.73% | 9.01% | 8.85% | 8.48% |
| CTLA-4$^{+}$LAG-3$^{-}$PD-1$^{-}$TIGIT$^{+}$TIM-3$^{-}$ | 1.18% | 1.48% | 0.63% | 4.27% | 4.72% | 1.67% |
| CTLA-4$^{+}$LAG-3$^{-}$PD-1$^{-}$TIGIT$^{-}$TIM-3$^{+}$ | 0.14% | 0.27% | 0.00% | 0.87% | 1.81% | 0.26% |
| CTLA-4$^{-}$LAG-3$^{+}$PD-1$^{+}$TIGIT$^{-}$TIM-3$^{-}$ | 10.60% | 14.48% | 3.96% | 2.76% | 3.47% | 1.49% |
| CTLA-4$^{-}$LAG-3$^{+}$PD-1$^{-}$TIGIT$^{+}$TIM-3$^{-}$ | 1.09% | 1.85% | 0.00% | 2.11% | 2.46% | 0.77% |
| CTLA-4$^{-}$LAG-3$^{+}$PD-1$^{-}$TIGIT$^{-}$TIM-3$^{+}$ | 3.14% | 5.59% | 0.58% | 0.26% | 0.42% | 0.13% |
| CTLA-4$^{-}$LAG-3$^{-}$PD-1$^{+}$TIGIT$^{+}$TIM-3$^{-}$ | 7.70% | 7.96% | 5.99% | 9.62% | 7.08% | 9.10% |
| CTLA-4$^{-}$LAG-3$^{-}$PD-1$^{+}$TIGIT$^{-}$TIM-3$^{+}$ | 1.04% | 1.33% | 0.56% | 1.14% | 1.51% | 0.68% |
| CTLA-4$^{-}$LAG-3$^{-}$PD-1$^{-}$TIGIT$^{+}$TIM-3$^{+}$ | 0.14% | 0.30% | 0.00% | 0.77% | 1.85% | 0.00% |
| **3 Checkpoints** | | | | | | |
| CTLA-4$^{+}$LAG-3$^{+}$PD-1$^{+}$TIGIT$^{-}$TIM-3$^{-}$ | 3.36% | 5.20% | 1.34% | 5.52% | 4.41% | 5.20% |
| CTLA-4$^{+}$LAG-3$^{+}$PD-1$^{-}$TIGIT$^{+}$TIM-3$^{-}$ | 1.56% | 3.03% | 0.00% | 4.47% | 6.29% | 1.49% |
| CTLA-4$^{+}$LAG-3$^{+}$PD-1$^{-}$TIGIT$^{-}$TIM-3$^{+}$ | 0.84% | 1.90% | 0.00% | 0.27% | 0.52% | 0.00% |
| CTLA-4$^{+}$LAG-3$^{-}$PD-1$^{+}$TIGIT$^{+}$TIM-3$^{-}$ | 9.61% | 12.78% | 4.46% | 9.94% | 7.57% | 10.40% |
| CTLA-4$^{+}$LAG-3$^{-}$PD-1$^{+}$TIGIT$^{-}$TIM-3$^{+}$ | 1.05% | 1.47% | 0.30% | 0.83% | 0.90% | 0.84% |
| CTLA-4$^{+}$LAG-3$^{-}$PD-1$^{-}$TIGIT$^{+}$TIM-3$^{+}$ | 0.46% | 0.66% | 0.21% | 0.32% | 0.56% | 0.00% |
| CTLA-4$^{-}$LAG-3$^{+}$PD-1$^{+}$TIGIT$^{+}$TIM-3$^{-}$ | 7.26% | 5.67% | 7.55% | 5.24% | 6.10% | 4.13% |
| CTLA-4$^{-}$LAG-3$^{+}$PD-1$^{+}$TIGIT$^{-}$TIM-3$^{+}$ | 5.00% | 10.75% | 0.11% | 0.71% | 0.85% | 0.32% |
| CTLA-4$^{-}$LAG-3$^{+}$PD-1$^{-}$TIGIT$^{+}$TIM-3$^{+}$ | 0.36% | 0.67% | 0.00% | 0.30% | 0.61% | 0.00% |
| CTLA-4$^{-}$LAG-3$^{-}$PD-1$^{+}$TIGIT$^{+}$TIM-3$^{+}$ | 1.12% | 1.60% | 0.00% | 2.94% | 3.47% | 1.93% |
| **4 Checkpoints** | | | | | | |
| CTLA-4$^{+}$LAG-3$^{+}$PD-1$^{+}$TIGIT$^{+}$TIM-3$^{-}$ | 8.10% | 4.31% | 7.69% | 9.38% | 7.19% | 6.04% |
| CTLA-4$^{+}$LAG-3$^{+}$PD-1$^{+}$TIGIT$^{-}$TIM-3$^{+}$ | 0.76% | 1.48% | 0.00% | 1.35% | 1.34% | 1.34% |
| CTLA-4$^{+}$LAG-3$^{+}$PD-1$^{-}$TIGIT$^{+}$TIM-3$^{+}$ | 0.64% | 1.32% | 0.00% | 1.69% | 2.87% | 0.42% |

| | | | | | | |
|---|---|---|---|---|---|---|
| CTLA-4$^{+}$LAG-3$^{-}$PD-1$^{+}$TIGIT$^{+}$TIM-3$^{+}$ | 1.42% | 1.41% | 1.38% | 2.48% | 2.87% | 2.23% |
| CTLA-4$^{-}$LAG-3$^{+}$PD-1$^{+}$TIGIT$^{+}$TIM-3$^{+}$ | 2.89% | 3.77% | 0.84% | 2.23% | 3.77% | 0.96% |
| **5 Checkpoints** | | | | | | |
| CTLA-4$^{+}$LAG-3$^{+}$PD-1$^{+}$TIGIT$^{+}$TIM-3$^{+}$ | 4.93% | 3.57% | 4.98% | 3.63% | 5.96% | 1.85% |

#### MATERIALS AND METHODS

##### Patients Demographics and Tissue Isolation

Fresh primary tumor tissue was obtained from patients undergoing surgical resection for histologically confirmed pancreatic ductal adenocarcinoma (PDAC) via a Whipple at the Winship Cancer Institute of Emory University (Atlanta, GA) following informed consent. All patients underwent surgery between 2019 and 2024. Freshly isolated liver punch biopsies of metastatic tissue were obtained from patients with histologically confirmed, inoperable, metastatic PDAC enrolled in the therapeutic clinical trial NCT03095781/WCI3321-16 and NCT04191421/WCI4463. Patients were excluded based on previous treatment with α-PD-1 and/or α-PD-L1. All patients in this study were enrolled between 2017 and 2022 and completed chemotherapy or other second-line regimens at least two weeks prior to biopsy and were off-treatment at time of sample acquisition. Details on patient demographics are reported in **Supplemental Tables 1 and 2**. Where possible, sex, race, ethnicity and age were accounted for to accurately represent the larger population of patients. Both male and female patients were included in each sample group.

##### Tissue Preservation and Quality

Samples were placed in cassettes and fixed for 24-48 hours in 10% neutral buffered formalin (NBF) prior to processing and embedding in paraffin. FFPE tissue blocks were sectioned at a thickness of 4 µm for all H&E, IHC and mIHC stains. H&E-stained tissues were analyzed by a pathologist to confirm presence of tumor and validate quality for further histological analysis. Samples that did not meet pathological requirements were excluded.

##### Immunohistochemistry and Multiplex Immunohistochemistry

Immunohistochemistry (IHC) and Multiplex immunohistochemistry (mIHC) was performed using automated protocols on the DISCOVERY ULTRA autostainer (Roche Diagnostics). Briefly, following deparaffinization, heat-induced epitope retrieval (HIER) was performed at 95°C using Cell Conditioning 1 (CC1, Roche, # 06414575001) for 64 min. Application of DISCOVERY Inhibitor (Roche, # 7017944001) was used to block endogenous peroxidase activity prior to antigen staining. Antigen staining was accomplished by an iterative incubation process with six primary antibodies. Briefly, each antibody was incubated for 40 min followed by a 12 min incubation with the corresponding OmniMap HRP secondary antibody (Roche, α-mouse # 05269652001 or α-rabbit # 05269679001) and then a 16 min incubation with a Opal™ tyramide signal amplification fluorophore (Akoya Biosystems, **Supplemental Table 5**) diluted using 1X Amplification Diluent (Akoya Biosystems, #FP1609) or Disc. Diluent P.S.S. for Opal 780 only (Roche, # 05266815001) and lastly followed by a denaturation cycle for 8 min at 93°C using Cell Conditioner 2 (CC2, Roche, # 05279798001). Tissue was stained with individual (IHC) or a combination of (mIHC) antibodies directed against the following markers: αSMA, CD3, CD4, CD8, CD19, CD68, CK19, FOXp3, IL-6, RORγt and T-bet (**Supplemental Table 5**). Where applicable, primary antibodies were diluted using Diamond: Antibody Diluent (Cell Marque, # 938B-09). Slides were counterstained with Spectral DAPI (Akoya Biosystems, # FP1490) for 16 min. Upon completion of the staining process and application of liquid cover slips (Mercedes Medical, # MER R2450), slides were removed from the autostainer and briefly soaked in a detergent solution to remove liquid coverslip residue and then glass cover-slipped using Vectashield® Antifade mounting medium (Vector Laboratories, # H-1000-10). Slides were cured for 24 hr in the dark and stored at 4°C prior to imaging. The Vectra® Polaris™ Automated Quantitative Pathology Imaging System was used to acquire whole slide multispectral images. Ideal exposures were determined by averaging the exposures generated by the Vectra Polaris software across a range of slides.

##### Analysis of mIHC Data

Whole-slide scans from the fluorescent-labeled sections were analyzed using QuPath v0.4.4.4 (University of Edinburgh. Edinburgh, Scotland, UK)[1]. DAPI channel (C1) was used for cell segmentation using the following settings: 0.4967 µm pixel size, 6.0 µm background radius, background by reconstruction, 1.0 µm median filter radius, 0.5 µm sigma size, 3.0 $µm^2$ minimum area, 400 $µm^2$ maximum area, threshold 5, 4.0 µm cell expansion area. After cell detection, cells were classified by adapting a publicly available script (https://www.imagescientist.com/creating-a-classifier) with threshold values for each fluorescent channel determined by defined visual cutoffs (fluorescent intensity ratio) to determine the number of positive cells for each fluorescent signal intensity or combination of signal intensities. All whole-image sections were analyzed simultaneously via the script editor in batch mode. Data was exported and analyzed using Microsoft Excel (V. 16.93) and GraphPad Prism (V. 10.4) tools. Percentages were calculated based on the total cell detection for each florescent signal or combination of signals divided by the total number of cells. Cellular density was calculated by dividing the total cell detection for each florescent signal, or combination of signals, by the total area of tissue.

**Single cell spatial analysis**

Whole tissue scans underwent quality control in QuPath prior to exporting of single cell metadata as follows: tissue that contained fractured pieces were separated and the largest intact piece of tissue was chosen for downstream analysis, excluding tissue not suitable for spatial analysis. Cell neighborhoods, cell-to-cell interactions, cell colocalization, spatial heterogeneity, definition of tumor margins and quantification of non-cancer cell populations relative to tumor structures was analyzed using the base object SPIAT from the SpatialExperiment[2]. SPIAT was used in R version 4.2 and tissue section mapping was used in Python version 3.9.18. Analysis included evaluating parameters of each individual tissue to assess tissue heterogeneity, as well as combining data of primary, or metastases tissue for comparison between tissues. Marker intensities from exported Qupath data were treated like gene expression levels, and cell coordinates and phenotypes as additional metadata. Using the spinglass algorithm from the igraph package in R with a set seed of 123 to ensure reproducibility, cell communities we delineated[3]. Briefly, network nodes (representing percent of cell type) and edges (representing the minimum distance between cell types) was used to detect cellular communities by optimizing the modularity of the network and organizing the cells into distinct groups based on their connectivity patterns. Each identified community is visualized with a background bubble situated behind network nodes and edges, indicating the community to which corresponding cells belong and allowed for the systematic categorization of cell types into closely associated groups based on their spatial distribution. Analysis of variables considered that most functions are independent of each other. Tissues with clear boundaries separating tumoral and stromal regions were included for infiltration analysis.

**Tissue Digestion**

All fresh tissue was subjected to a commercial mechanical/enzymatic dissociation system (GentleMACS, Miltenyi Biotec, Bergish Gladbach, Germany). GentleMACS dissociation was performed according to the manufacturer’s protocol for tough tumors. Briefly, the tissue was cut into small fragments about 2–4 mm in length and put in a C-tube (Miltenyi Biotech) with RPMI 1640 and enzymes H, R and A (Miltenyi Biotec) according to the manufacturer’s recommendation. This mixture was then subjected to two cycles of mechanical dissociation using program h_tumor_01 (program repeated two times) with the GentleMACS dissociator, followed by a 30 min incubation at 37 °C while shaking. Tumor mixture was then subjected to an additional round of mechanical dissociation and single cell suspensions were generated by filtration through a 70 µm filter. Filtered cell suspension was centrifuged at 1700 rpm for 5 min. Cell pellets, resuspended in 1X PBS, containing $2x10^6$ cells or less per sample were then processed for CyTOF staining.

**Single-Cell Mass Cytometry Staining**

Viability staining was conducted by incubating cells with 1X PBS and Cell-ID™ Cisplatin (Fluidigm Sciences). Staining was quenched using Maxpar® Cell Staining Buffer and washed twice before surface and intracellular staining following manufacturer's protocol (Fluidigm Sciences). Cells were stained with the following surface markers: CD45-89Y, B7H3-141Pr, CD19-142Nd, HLADR-143Nd, CD69-144Nd, CD4-145Nd, CD8-146Nd, ICOS-148Nd, CD127-149Sm, CD103- 151Eu, CD95-152Sm, TIGIT-153Eu, TIM3-154Sm, CD27-155Gd, PDL1-156Gd, CD33-158Gd, CCR7-159Tb, CXCR5-164Dy, CD45RO-165Ho, NKG2D-166Er, CD11b-167Er, CD25-169Tm, CD3-170Er, CD38-172Yb, CXCR4-173Yb, PD1-174Yb, CD14-175Lu, CD56-176Yb and CD16-209Bi (**Supplemental Table 9**). Following surface staining, cells were washed, fixed, permeabilized, and stained with the following intracellular markers: TCF1-147Sm, LAG3-150Nd, TBet-160Gd, CTLA4-161Dy, FoxP3-162Dy, EOMES-163Dy, Ki67-168Er and Granzyme B-171Yb. Following intracellular staining, cells were washed and stained with Cell-ID™ Intercalator-Ir 125μm (Fluidigm Sciences). Samples were acquired on Helios mass cytometer (Fluidigm Sciences). Acquired raw FCS files were normalized with the preloaded normalizer algorithm on CyTOF software.

**Single-Cell Mass Cytometry Data Analysis**

Normalized CyTOF FCS files were analyzed using both Flowjo (V. 10.10) and Cytobank (V. 10.6) software[4]. FSC files were manually cleaned using Fluidigm technical notes (PN 400248 B1) cleanup strategy. Briefly, normalization beads were removed from samples by gating on the negative population. A series of Gaussian parameters (Residual, Center, Offset, and Width) were used to eliminate events from merged ion clouds. Event Length was used to filter out abnormal signal pulses. Live cells were identified and gated as cisplatin negative populations. Iridium intercalator gates labeled DNA1 and DNA2 identified intact singlet nucleated cells. Samples containing at least 500 live cells ($DNA2^+$) were included for further analysis (primary, n = 12; metastatic, n = 23). Detailed gating strategies for all analyses can be found in Figure S11 – S15. Cleaned FCS files were exported from Flowjo as individual sample files. Files were uploaded into Cytobank for high dimensional clustering analysis. $DNA2^+$ cells were subjected to optimized parameters for T-distributed stochastic neighbor embedding (opt-SNE) analysis[5], with equal event sampling selected to ensure samples containing more live cells did not skew clustering. Files were analyzed using default perplexity (30) and iterations (1000) using 35 markers and seed 1845947262. Opt-SNE plots were displayed as either individual sample or concatenated samples based on primary or metastatic grouping using Cytobank Illustration Editor. Healthy donor peripheral blood mononuclear cells (PBMCs) were used to inform manual gates.

**Statistical Analysis**

Statistical significance was analyzed with GraphPad Prism (V. 10.4) software. For all comparisons, $p$-value < 0.05 was considered statistically significant. Comparisons between two experimental groups were performed using unpaired two-tailed Mann–Whitney test (for data not normally distributed) or Wilcoxon t-test (for matched patient samples). For multiple comparisons, two-way ANOVA with post hoc Sidak's or Bonferroni's analysis was performed. Statistical details of individual experiments are found in the corresponding figure legends.

**Data Availability**

All data reported in this paper will be shared by the corresponding author upon reasonable request.
All original code is available in this paper's supplemental information.
Any inquiries regarding data access or analyses reported in this paper should be directed to the the corresponding author upon request.
